# Therapeutically targeting the classical complement pathway with antisense oligonucleotides in Alzheimer’s disease

**DOI:** 10.1101/2024.08.23.609240

**Authors:** Yuanyuan Wang, Tiffany Wu, Alvin Gogineni, Ming-Chi Tsai, Lukasz Kielpinski, Anubha Mahajan, Shristi Pandey, Luis Schachner, John C. Tran, Cong Wu, Stephen Schauer, Anne Biever, Sidsel Boesen, Meiling Li, Alexander Munk, William J Meilandt, Justin Elstrott, Amy Easton, Helene M Gylling, Jesse E Hanson

**Affiliations:** Department of Neuroscience, Genentech Inc., South San Francisco, CA 94080, USA; Department of Translational Imaging, Genentech Inc., South San Francisco, CA 94080, USA; Roche Pharma Research and Early Development, Therapeutic Modalities, Roche Innovation Center Copenhagen, Hørsholm DK 2970; Department of Human Genetics, Genentech Inc., South San Francisco, CA 94080, USA; Department of OMNI Bioinformatics, Genentech Inc., South San Francisco, CA, 94080 USA; Department of Biochemical and Cellular Pharmacology, Genentech Inc., South San Francisco, CA 94080, USA; Department of Translational Medicine, OMNI, Genentech Inc., South San Francisco, CA, 94080 USA; Pharma Research and Early Development, Therapeutic Modalities, Roche Innovation Center Basel, F. Hoffmann-La Roche Ltd., 4070 Basel, Switzerland

**Author notes:** These authors contributed equally (Yuanyuan Wang, Tiffany Wu, Alvin Gogineni, Ming-Chi Tsai).

## Abstract

The complement classical pathway (CP) is a key mediator of synapse loss and neurodegeneration in mouse models of Alzheimer’s (AD) and other neurodegenerative diseases. We analyzed human brain proteomics and found consistent elevations of all CP proteins, but not other complement pathways, in AD patient brains. We performed human genetics analysis that identified a rare variant in the *C1S* gene within the Finnish population that is associated with AD and we found that a common AD-associated *C1S* variant correlates with increased C1S protein levels. A targeted assay detected elevated C1S activation in AD patient CSF. Given this specific implication of the CP in AD, we next evaluated the therapeutic approach of targeting the CP in the brain using antisense oligonucleotides (ASOs). To identify promising CP targets for knockdown using ASOs we first tested for rescue of synapse loss in an AD mouse model using heterozygous and homozygous complement knockout mice and examined the relative brain expression levels of different CP genes. Based on these experiments we prioritized C1r, C1s and C4 as promising targets for therapeutic knockdown using ASOs. We then screened for ASOs for each target, evaluating *in vitro* and *in vivo* knockdown and toxicity, and identified optimal ASOs targeting C1r, C1s and C4. Experiments with AD model mice demonstrated significant rescue of synapse loss following treatment with C1r, C1s or C4 ASOs. Overall, our findings provide proof of concept for using nucleic acid-based medicine to target the CP in AD and demonstrate the translational potential of this approach.

## Introduction

In addition to functioning in humoral innate immunity, the complement classical pathway (CP) contributes to microglial-mediated synapse removal in the brain during early postnatal neural circuit refinement (*1–3*) (Fig. 1A). This process involves synapse tagging by the C1 complex (consisting of C1q, C1r, and C1s). C1q binding activates the C1r protease, which cleaves and activates the C1s protease. Active C1s cleaves soluble complement components C4 and C2, resulting in the covalent attachment of the C3 convertase (C4bC2b) to the synapse. The C3 convertase then cleaves soluble C3, resulting in the attachment of the C3b fragment to the synapse. Microglia detect synapses tagged by C3b via CR3 receptors resulting in synapse elimination (*4, 5*). While physiological CP function in synapse removal is restricted to specific circuits during early postnatal development (e.g. retinal ganglion cell synapses in the lateral geniculate nucleus), recent research has identified widespread and detrimental effects of CP activation in the context of neurodegenerative disease (*4, 6, 7*). In particular several studies have shown complement plays a role in amyloid and tauopathy models of Alzheimer’s disease (*8–13*). The sensitive nature of CP regulation in the brain is highlighted by recent work showing that the secreted neuronal pentraxin NPTX2 can regulate CP activity by binding to C1q (*13*). Loss of NPTX2 results in complement-dependent synapse loss, while overexpression protects synapses in a tauopathy model (*13*). Complement gene expression is induced by activated glial cells in response to many pathologies, and accordingly the complement CP has been shown to play a detrimental role in various neurodegeneration models in addition to AD. This includes models of frontotemporal dementia (FTD), Huntington’s disease, and virus-induced memory impairment (*14–16*). Furthermore, GWAS studies have found that haplotypes with increased C4 expression levels are associated with schizophrenia risk (*17, 18*), as are variants in the neuronal CP inhibitor CSMD1 (*19–21*).

**Fig. 1.**
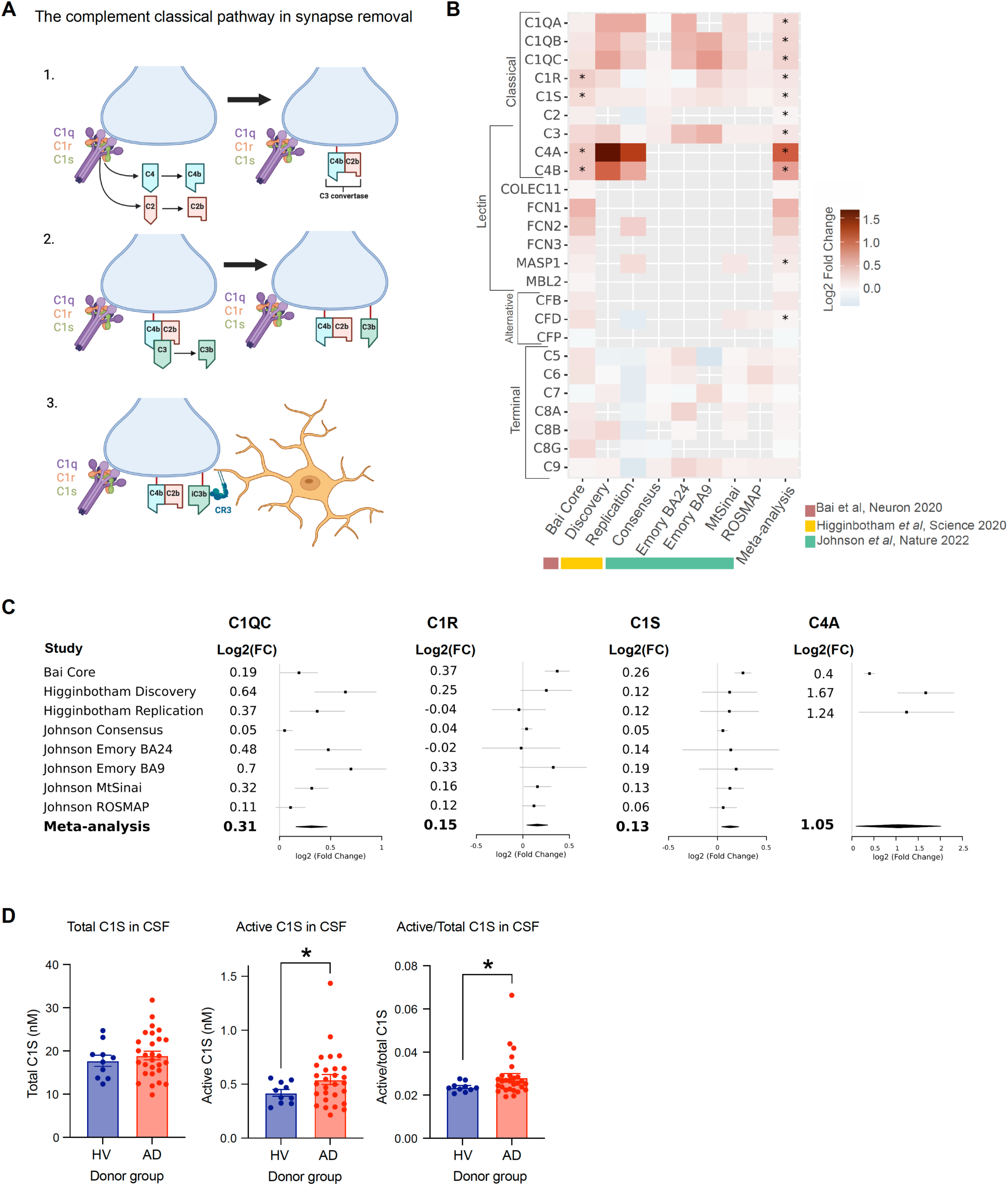
Specific elevation and activation of the classical pathway in AD patients. (**A**) Schematic diagram showing the complement classical pathway function in synapse tagging for glial-mediated synapse elimination (created with BioRender.com). C1 complex binding to a synapse leads to deposition of C4b and C2b fragments (1) which is followed by C3b deposition (2) and recognition C3b by microglia (3). (**B**) Meta-analysis of complement protein brain expression: Heatmap of all complement proteins. Brain tissue proteomics data from 8 different cohorts including AD and control samples was analyzed from Bai et al (*24*), Higginbotham et al (*25*) and Johnson et al (*26*). Heatmap shows log fold changes in protein abundance for each dataset in AD vs control as well as a meta-analysis of all datasets (see methods). Only datasets where proteins were detected were used to combine fold changes and p-values in the meta-analysis. Asterisks indicate significant log fold change differences in individual data sets (limma) or in the meta-analysis (Fisher’s method) as described in the methods section. Empty slot in the heatmap reflects that the give protein was not detected in that dataset. (**C**) Fold changes and forest plots showing effect sizes and 95% confidence intervals in the individual data sets and meta-analysis for example classical pathway proteins (C1QC, C1S, C1R, C4A). (**D**) Plots show measurements of levels of total C1S and active C1S, and the ratio of the two measures in human CSF from Alzheimer’s disease patients (AD, n = 28) and healthy volunteers (HV, n = 10) using liquid chromatography mass spectrometry (LC-MS) analysis of surrogate peptides (active C1S peptide: IIGGSDADIK; total C1S peptide: LLEVPEGR). LC-MS quantitation of C1S levels was calculated using a signal calibration equation determined with a heavy-labeled peptide standard. Demographics of healthy and Alzheimer’s disease cerebrospinal fluid donors are listed in Table 1. Data are shown as mean +/- SEM. *, p < 0.05, Welch’s t-test for significance of the difference of the means.

**Table 1, related Fig. 1D.**
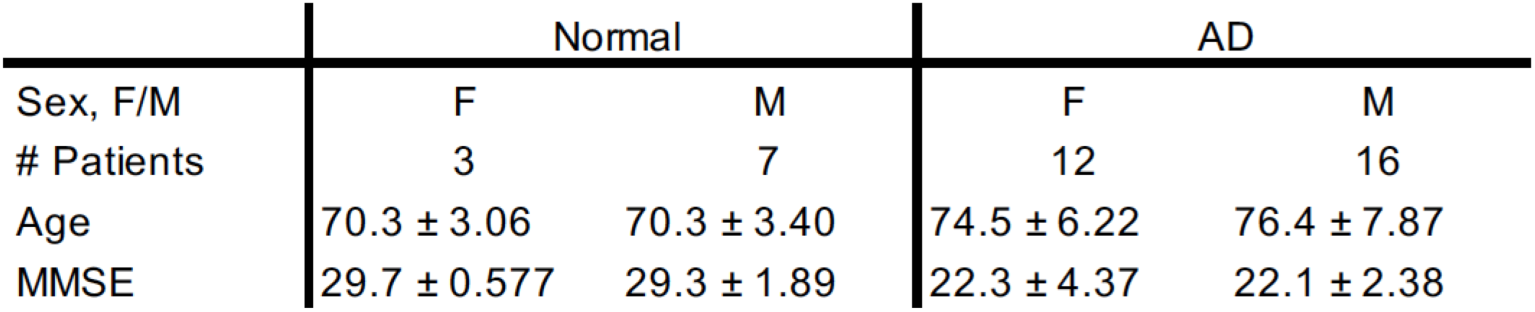
Demographics of healthy and Alzheimer’s disease cerebrospinal fluid donors.

While mouse AD models show a detrimental role for the CP, and measurements in AD patient CSF show elevations in intact and active C3 and C4 (which are downstream of other pathways as well as the CP) (*11, 12, 22*), we sought evidence for specific and causal roles of the CP in human AD. Here we describe new genetic analyses of the CP and analyses of CP protein levels in AD patient CNS samples. These analyses found associations of C1s with AD and specifically point to a role for excess CP activity in AD. Together these observations make the CP an attractive therapeutic target for CNS disorders. However, challenges come from the high levels of complement in systemic circulation. In particular, systemically administered therapeutics will have to overcome difficulties with achieving CNS target engagement as well as potential safety considerations related to suppressing peripheral innate immune function. In contrast, CNS-delivered anti-sense nucleotides (ASOs) have the potential advantages of direct, selective, and sustained inhibition of the CP in the brain (*23*). We therefore sought to identify ASO targets within the CP. Our previous work found that C3 KO can protect against plaque-associated synapse loss in the PS2APP model of amyloidosis (*11*). However, C3 KO blocks not only the CP, but also the lectin pathway (LP) and the alternative pathway (AP) of the complement system, and in addition to providing synapse protection, C3 KO also increases plaque load in amyloidosis model mice (*9, 11*). Therefore, we examined the effects of knocking out components of the CP upstream of C3 in the PS2APP model. To evaluate the therapeutic potential of ASOs, which are unlikely to achieve the level of target knockdown corresponding to homozygous KO, we included heterozygous KO mice in these experiments. Based on the results of these KO experiments and other considerations, including assessment of CP component brain expression levels, we prioritized C1r, C1s and C4 as potential ASO targets. We then pursued discovery of well tolerated and effective ASOs targeting each of these genes and used these ASOs in preclinical efficacy studies in PS2APP mice, which demonstrated significant beneficial effects of synapse rescue.

## Results

### Proteomics analyses show elevated CNS CP levels and activation in AD patients

To comprehensively assess expression of complement pathway components in the human brain, we performed a meta-analysis using published data from 8 different cohorts that had brain proteomics from AD patients and healthy controls (*24–26*). This analysis revealed significant elevations of all of the classical pathway proteins, while complement proteins involved specifically in the AP, LP, or terminal pathways were not generally elevated in AD patient brains (Fig. 1B,C). This is consistent with a role for elevated CP, but not AP, LP or terminal pathway production by activated brain cells, rather than a general infiltration of blood-derived complement proteins in AD patient brains (which would result in elevation of all complement proteins). In addition, the levels of some regulators and receptors for the complement pathway components were also altered in AD patient brains (Fig. S1).

We have previously applied targeted immunoassays and found elevated levels of activated C3 (which is downstream of the AP and LP as well as the CP) and activated C4 (which is downstream of the LP as well as the CP) in AD patient CSF samples (*11, 12, 22*). To more specifically assess CP activity, we have now developed liquid chromatography-mass spectrometry (LC-MS) assays for detecting total and active (processed) C1S in cerebrospinal fluid (CSF). Measurement of C1s levels in AD patient and healthy volunteer CSF samples showed significant elevation of active C1S levels as well as the ratio of active:total C1S in AD patient CSF (Fig. 1D; Fig. S2). This specifically indicates increased classical pathway activity in the CNS of AD patients.

### Human genetics analyses implicate *C1S* in AD

We next evaluated the potential causal relationship between the CP and AD susceptibility. The most recent AD GWAS (Stage 1) displayed a nearly genome-wide significant association in proximity to the *C1S* gene, where the index SNP is a common non-coding variant, rs3919533 (T allele, frequency 82%, p=9.9x10^-7^, odds ratio [95% CI]=1.06[1.03-1.09]), located 5Kb upstream of *C1S* (Fig. 2A)(*27*). To assess the impact of the variant on protein levels, we analyzed an o-link plasma proteomics dataset from the general population (UK Biobank samples) (*28*), which included data on 2,923 proteins and searched for proteins displaying changed levels between risk allele carriers and protective allele carriers. In this analysis, C1S was identified as the plasma protein with the most significant and highest fold increase in AD risk allele carriers (Fig. 2B), indicating that the impact of variant rs3919533 on AD risk is linked to elevated C1S levels.

**Fig. 2.**
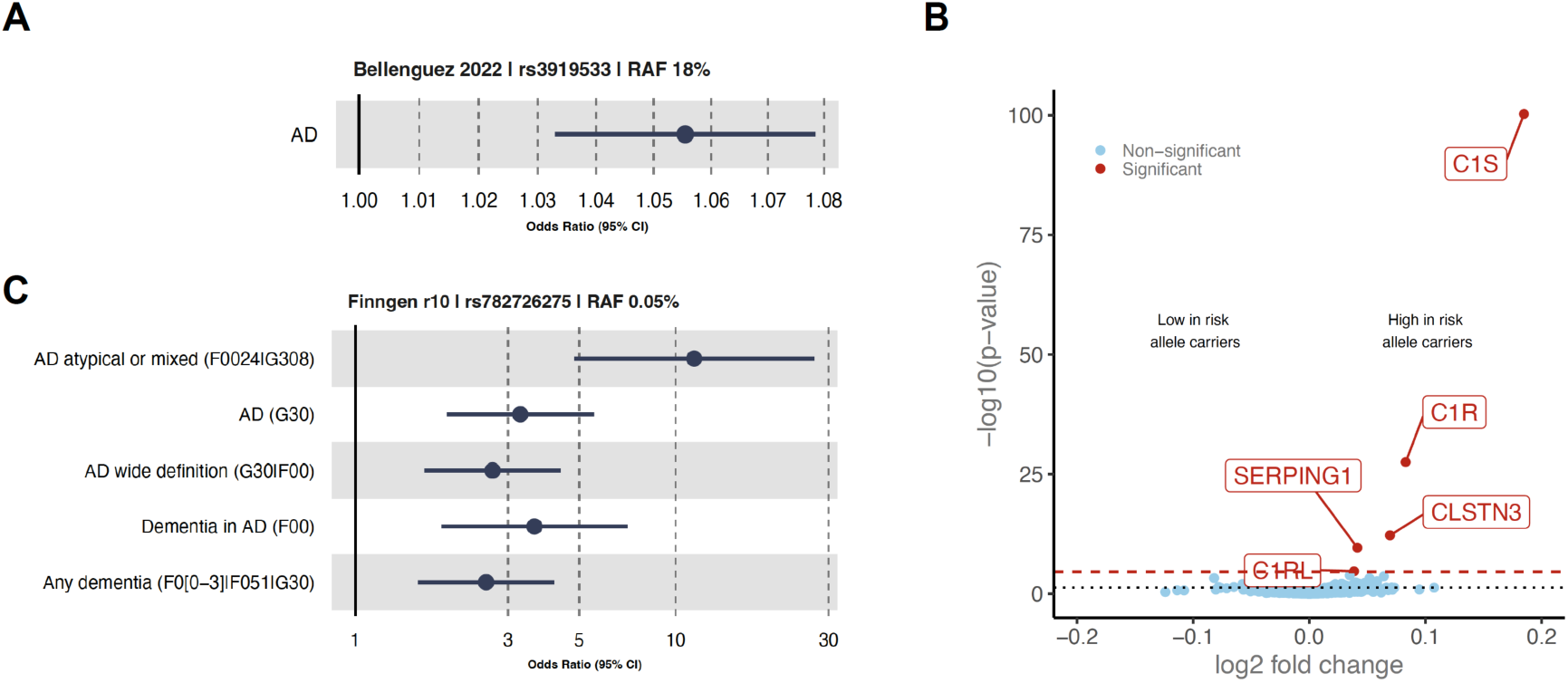
Human genetics implicate increased C1S in AD risk. (**A**) and (**C**) The forest plots represent the odds ratios, confidence intervals for the association of rs3919533 (**A**) and rs782726275 (**C**) with AD in various AD groups. The ICD10 codes used to define AD are indicated in parentheses. RAF stands for risk allele frequency and CI denotes confidence interval. P value and case and control numbers are listed in Table 2. (**B**) The volcano plot illustrates the differentially expressed proteins in the plasma of rs3919533 risk allele carriers compared to non-risk allele carriers. The red dashed line signifies the Benjamin-Hochberg threshold. Proteins that are differentially expressed (with false discovery rates less than 0.05) are emphasized in red.

**Table 2, related to Fig. 2A,C.** Healthy and Alzheimer’s disease donor sample numbers and odds ratio p-values for studies shown in Fig. 2A,C.

| Study | p-value | n_case | n_control |
| --- | --- | --- | --- |
| AD | 9.96E-07 | 85665 | 400902 |
| AD atypical or mixed (F0024\|G308) | 3.43E-08 | 2419 | 183753 |
| AD (G30) | 1.16E-05 | 10520 | 401661 |
| AD wide definition (G30\|F00) | 6.25E-05 | 15617 | 396564 |
| Dementia in AD (F00) | 1.64E-04 | 6145 | 388560 |
| Any dementia (F0[0-3]\|F051\|G30) | 2.07E-04 | 16209 | 301879 |

Additionally we examined data from the FinnGen study (release 10, https://r10.finngen.fi/)(29), which was not part of the recent AD GWAS stage 1, and identified an independent (European r2<0.2) rare Finnish-enriched (3X) intronic variant, rs782726275 (G allele, frequency 0.05%, p= 0.00021-3.4x10^-8^, odds ratio =2.5-11.4), within *C1S* that is associated with AD risk with varying strengths depending on the AD definition (Fig. 2C). Notably, the risk associated with this variant was particularly pronounced in the mixed phenotype of AD (Fig. 2C).

These observations of both rare and common variant associations, and the assessment of the common variant effects on protein levels, collectively demonstrate an association of *C1S* with AD risk and suggest that the risk is linked to elevated C1s levels. Overall, this evidence suggests that C1S might have a causal role in predisposing individuals to AD.

### Evaluating potential CP ASO targets

Given the important role of CP in human AD suggested by genetic and proteomic analyses of AD patients, we performed pre-clinical proof-of-concept studies to investigate the approach of knocking down CP genes in the brain using an AD mouse model. ASOs have been shown to efficiently knock down target gene expressions across cell types and regions of the brain, but do not achieve levels of target reduction comparable to homozygous deletion (*23, 30*). Therefore, we aimed to evaluate which components of the CP might be most sensitive for achieving inhibition of CP function in synapse loss with partial target reduction by ASOs. While the CP has been shown to play a role in both tauopathy and amyloidosis models for this study we focused on plaque associated-synapse loss in the PS2APP amyloidosis mice because we have found that this model shows robust complement-dependent synapse loss without requiring extensive aging of the mice (*11*). As our previous study used C3 KO, which blocks the CP, LP and AP, we here tested the effect of C1q KO, which only blocks the CP, in the same model. We first confirmed that C1q heterozygous and homozygous knockout mice expressed ∼50% WT levels, or no, C1q mRNA and protein respectively (Fig. S3A,B). Consistent with the previous study, we observed ∼50% reduction in spine density near amyloid plaques compared to spine density away from plaques (Fig. 3A,B). While heterozygous C1q KO did not significantly affect this plaque-associated synapse loss, homozygous C1q KO almost fully rescued the synapse loss (Fig. 3A,B). In contrast to previous results with C3 KO (*11*), C1q KO did not affect cortical Aβ plaque load in these PS2APP mice (Fig. 3C). To examine the CP downstream of C1q, we next assessed the role of C4. These experiments showed significant gene-dose dependent rescue of plaque-associated spine loss in C4 heterozygous and homozygous KO, PS2APP mice (Fig. 4A,B). We also confirmed that C4 heterozygous mice expressed half the amount of mRNA and protein compared with WT mice (Fig. S3C-E). Similar to C1qKO, C4 KO did not affect cortical amyloid plaque load in PS2APP mice (Fig. 4C). These results showing rescue with C1q or C4 KO support that complement-dependent synapse loss is mediated specifically by the CP. At the same time the lack of effects of C1q or C4 KO on amyloid plaque load support that the increased plaque load previously seen with C3 KO is CP-independent.

**Fig. 3.**
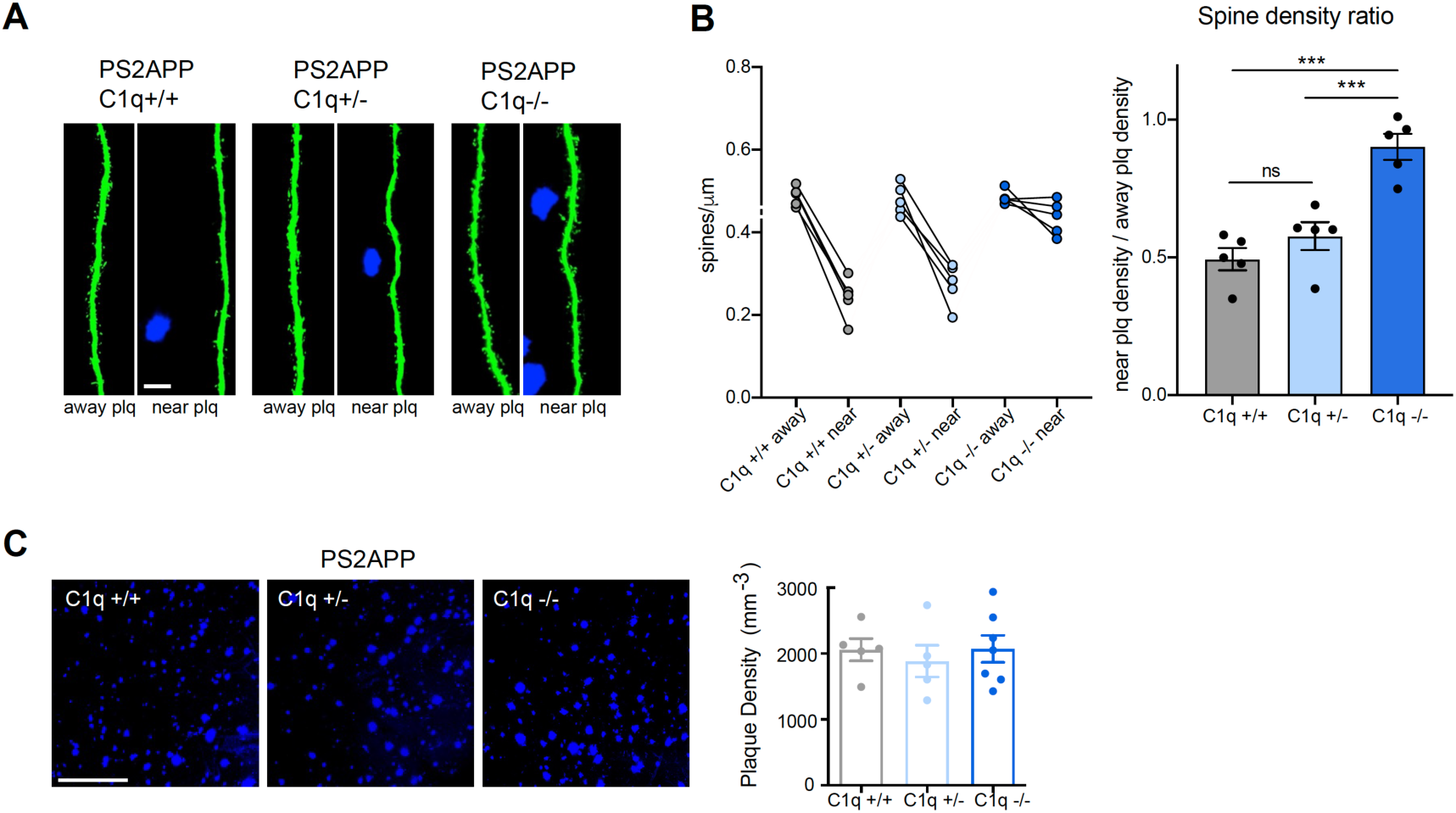
Synapse loss is rescued in C1q -/- mice, but not in C1q +/- mice. (**A**) Two-photon *ex vivo* brain imaging of 6-month-old male PS2APP mice crossed to C1q KO mice. Representative images show GFP-expressing neuronal dendrites that are near or away from Aβ plaques (plq) in somatosensory cortex with genotypes as indicated. Plaques were stained with methoxy-X04 (blue). Scale bar, 20 μm. (**B**) *Left*, quantification of spine density as shown in (A). Each dot represents one mouse (average of 6 dendrites). Average spine density away from plaques and near plaques from each mouse are shown as connected dots. *Right*, the spine density ratio is shown for each mouse (the average near plaque spine density divided by average away plaque spine density). (**C**) Representative images (*left*) and quantification (*right*) of Aβ plaques labeled by methoxy-X04 in mouse brains with genotypes as indicated. Each dot represents a mouse. Scale bar, 200 μm. Data are represented by mean +/- SEM. ***, p < 0.001, one-way ANOVA with Tukey’s multiple comparisons test.

**Fig. 4.**
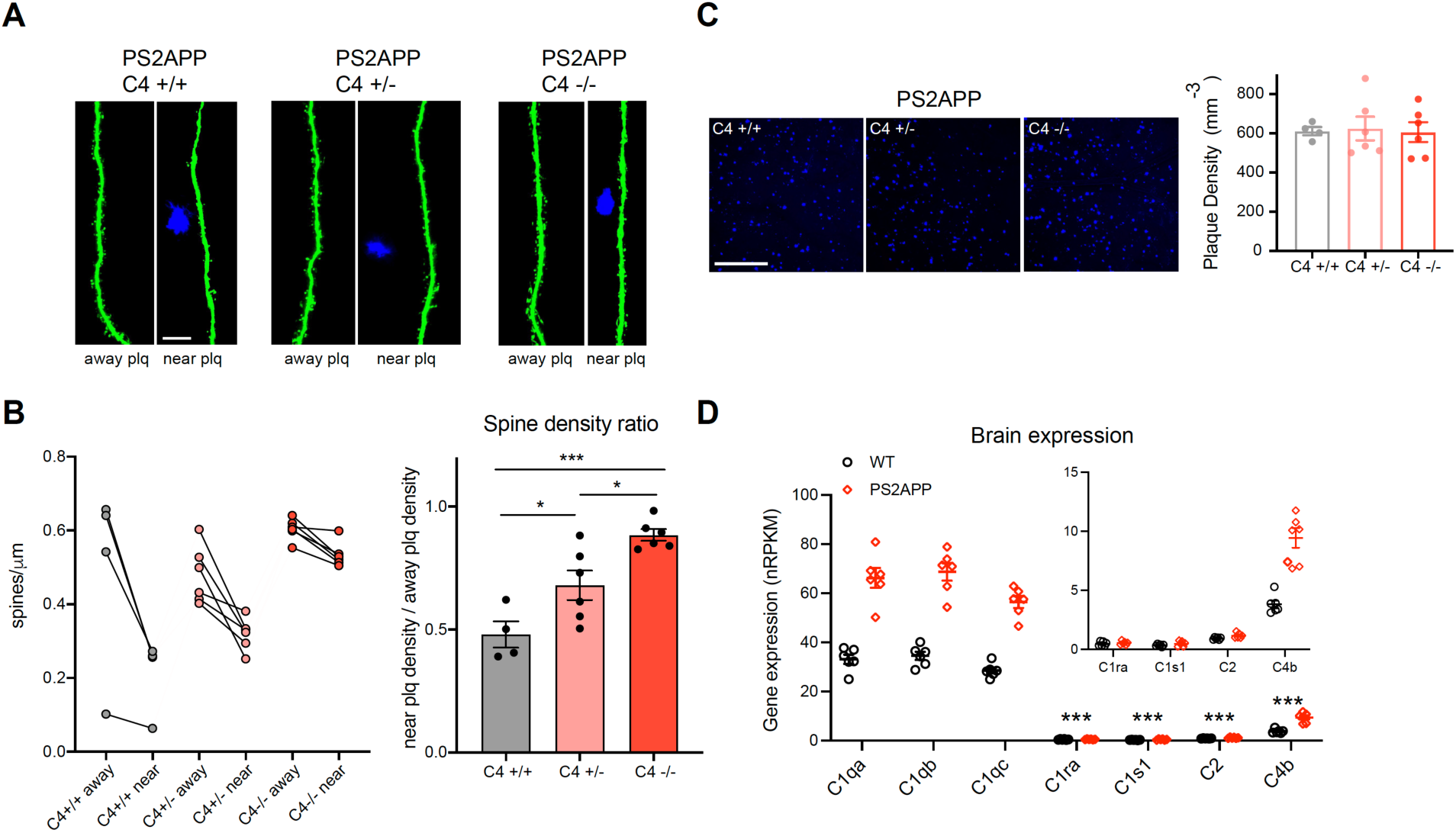
C4 -/- and C4 +/- mice show gene dose-dependent rescue of synapse loss. (**A**) Similar to Fig. 1, Two-photon *ex vivo* brain imaging 6-month-old male PS2APP mice crossed to C4 KO mice. Representative images show GFP-expressing neuronal dendrites that are near or away from Aβ plaques (plq) in somatosensory cortex with genotypes as indicated. Plaques were stained with methoxy-X04 (blue). Scale bar, 20 μm. (**B**) *Left*, quantification of spine density as shown in (A). Each dot represents one mouse (average of 6 dendrites). Average spine density away from plaques and near plaques from each mouse are shown as connected dots. *Right*, the spine density ratio is shown for each mouse (the average near plaque spine density divided by average away plaque spine density). (**C**) Representative images (*left*) and quantification (*right*) of Aβ plaques labeled by methoxy-X04 in mouse brains with genotypes as indicated. Each dot represents a mouse. Scale bar, 200 μm. Data are represented by mean +/- SEM. *, p < 0.05, ***, p < 0.001, one-way ANOVA with Tukey’s multiple comparisons test. (**D**) Classical pathway (C1q, C1r, C1s, C2, and C4) gene expression in WT and PS2APP mouse brain measured by bulk RNAseq (GSE75357). Inset shows C1r, C1s, C2 and C4 expression with zoomed in y-axis values. Data are represented by mean +/- SEM. C1r, C1s, C2 and C4 expression were statistically significantly lower compared with C1qa, C1qb, or C1qc expression, in either WT or PS2APP mice, ***, p < 0.001, one-way ANOVA with Dunnett’s multiple comparisons test.

The significant impact of even heterozygous C4 KO on synapse loss suggests C4 could be a sensitive target for protecting against synapse loss in disease. On the other hand, heterozygous C1q KO did not show significant protection of synapses. We next assessed brain expression levels of CP genes in mice. While C1q subunit genes (C1qa, C1qb, C1qc) are expressed at comparable high levels in mouse brain and liver, the other two components of the C1 complex, C1r and C1s, are expressed at much lower levels in the brain compared to liver (Fig. S4A). RNAseq analysis of brain tissues from WT and PS2APP mice found that while C1q is very abundant, C1r and C1s are expressed at much lower levels (Fig. 4D). It is therefore likely that C1q levels are in excess, while C1r and C1s levels may be rate-limiting in brain C1 complex formation. Given these considerations and our human genetics analyses implicating *C1S* in AD, we decided to focus on C1r and C1s as ASO targets for disrupting C1 function. While C4 and C2 were both also expressed at lower levels compared to C1q, we decided to prioritize C4 for additional ASO discovery based on our AD model efficacy data with C4 KO mice, as well as the genetic implication of *C4A* in schizophrenia.

### *In vitro* screens for C1r, C1s and C4 ASOs

We designed locked nucleic acid (LNA)-gapmer ASOs targeting mouse C1r, C1s and C4, taking into consideration factors including sequence homology across the duplicated genes in mouse (e.g. C1s1 and C1s2; C1ra and C1rb; C4b and C4a), ASO length, predicted hybridization free energy, and number of predicted mis-match off targets. Based on these considerations, we designed and synthesized approximately 700 ASO compounds in total for the three targets. The knockdown (KD) efficiency of these ASOs was first evaluated by RT-qPCR using mouse primary hepatocytes, which endogenously express high levels of complement proteins, allowing for easy target detection and sensitive screening. After the primary screen about 150 ASO with good target knockdown were selected for further safety screening using a caspase activation assay to detect toxicity (Fig. 5A). 49 compounds that passed the caspase activity screen, consisting of 23 C4 targeting ASOs, 16 C1s targeting ASOs and 10 C1r targeting ASOs, were then selected for secondary screening using mouse primary astrocytes. As C1ra, C1s1 and C4b are the higher expressed isoforms in mouse brains for the respective target genes (Fig. S4B), their expression was measured by qPCR in this screen. Four-point dose-response curves were generated for each ASO to calculate IC50 values in astrocytes (Fig. 5B). Comparison between the KD of C1r, C1s or C4 by ASOs in astrocytes and hepatocytes showed that the KD efficiency was well correlated in our primary and secondary screens (Fig. S5).

**Fig. 5.**
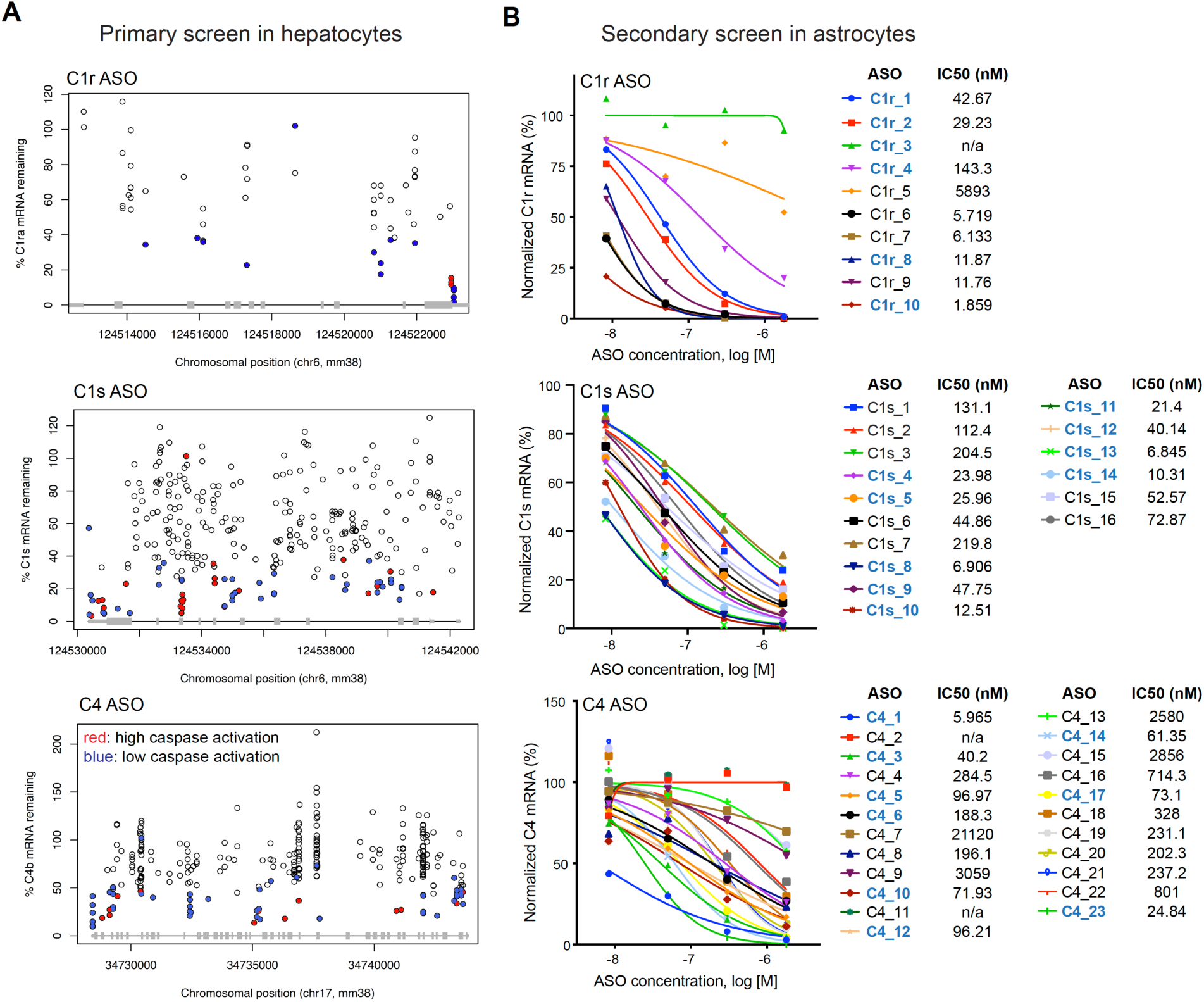
In vitro screening for ASOs against C1r, C1s and C4 identifies candidate ASOs with potent gene KD. (**A**) Quantification of residual target (C1ra, C1s or C4b as indicated) mRNA expression level after ASO (0.3 μM) incubation for 3 days in mouse primary hepatocytes measured by RT-qPCR. Target gene expression was first normalized to the housekeeping gene Rplp0 and then normalized relative to expression with saline treatment. Each dot represents one ASO. X-axis illustrates the binding position on the transcript of each gene. Selected ASOs with good knockdown efficiency were tested for toxicity using a caspase activation assay. ASOs that had low caspase activation are labeled in blue and ASOs that induced high caspase 3 activation are labeled in red. (**B**) Dose-response curves of knockdown of C1r, C1s or C4 (as indicated) by respective ASOs in mouse primary astrocytes assessed by RT-qPCR after 3-day ASO treatments. Target gene expression was normalized to expression in saline treated astrocytes after normalization to the housekeeping gene GAPDH. IC50s calculated from dose response curves are shown next to each ASO. ASOs indicated by blue bold font were selected for *in vivo* studies.

### *In vivo* characterization of CP ASOs

Nine C4 ASOs, nine C1s ASOs, and six C1r ASOs that exhibited good KD *in vitro* (Fig. 5B; blue-colored ASOs) were selected for evaluating tolerability and KD efficiency *in vivo* (Listed in Table 3). Targeting ASOs, a non-targeting control ASO (NT) (the last ASO in Table 1 of reference (*31*)), or a saline control were delivered via intracerebroventricular (ICV) injection (100 μg) and acute toxicity (within two hours) was assessed based on behavioral scoring (Fig. 6A-C). Only ASOs without significant acute toxicity were considered for further evaluation. Mouse brain tissues were collected after two weeks and brain C1r, C1s or C4 levels were measured by qPCR. Consistent with our in vitro screening data, many ASOs produced more than 50% target KD, and a few ASOs reached >80% target KD (Fig. 6A-C). In order to further evaluate potential toxicity, we measured the expression of 3 innate immune response markers that are expressed by glial cells and induced during neuroinflammation (Cst7, Gpnmb, Oasl2)(*32, 33*) (Fig. S6). ASOs that exhibited significant upregulation of these markers and/or resulted in body weight decline during the two-week period were excluded (Fig. 6D and Fig. S6). Based on favorable KD and tolerability, two C1r ASOs, three C1s ASOs, and three C4 ASOs were selected for further evaluation (Fig. 6A-C).

**Fig. 6.**
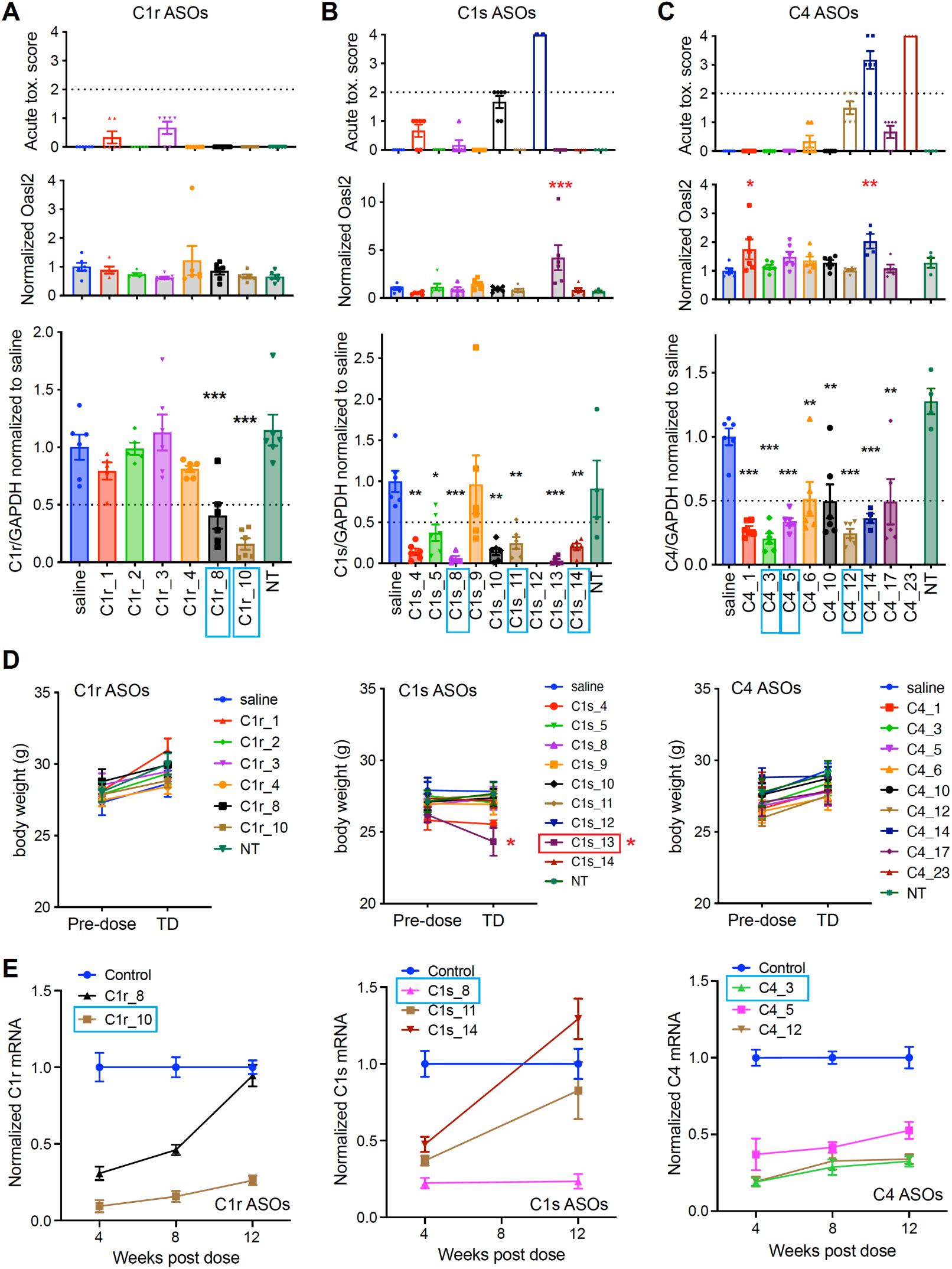
*In vivo* screening for ASOs against C1r, C1s and C4 identifies optimal ASOs for use in efficacy studies. (A) *Top*, C1r ASO induced toxicity after ICV injection was assessed by behavioral scoring (see methods). Only ASOs that induced score less than 2 were considered for further evaluation. *Middle*, expression of the inflammatory marker gene Oasl2 was measured by RT-qPCR from mouse brain tissues collected 14 days after ICV injection of ASOs. Data are normalized to Oasl2 expression in saline injected mice. *Bottom*, target gene C1r expression assessed by RT-qPCR. Data are normalized to C1r expression in saline injected mice. (**B**) and (**C**), similar to (A) with ICV injection of C1s ASOs (B) or C4 ASOs (C). Each dot represents a mouse, n = 4-6 mice per condition. Data are represented by mean +/- SEM. *, p < 0.05, **, p < 0.01, ***, p < 0.001, one-way ANOVA with Dunnett’s multiple comparisons test; compared with saline condition. ASOs in blue-colored boxes are the ASOs selected for in-vivo time course studies shown in (E). (**D**) Mouse body weights before and 14-day after saline/ASO injection. Plots show average mouse body weights per condition +/- SEM, n = 4-6 mice per condition. Only C1s_13 (red-colored box) induced trend of body weight decline (p = 0.05, student’s paired t-test, two-tail). (**E**) C1r, C1s or C4 mRNA expression (assessed by RT-qPCR) in the mouse brain 4 weeks, 8 weeks and 12 weeks after ICV injection of saline or ASOs. Control condition includes combined results from saline injected and non-targeting ASO injected mice. Target gene mRNA expression levels were normalized to the control condition at each time point after normalization to GAPDH expression. Plots show mean +/- SEM expression of the indicated genes in the mouse brains at each time point, n = 5-6 mice per group per time point. ASOs in blue-colored boxes were selected for in-vivo efficacy studies.

**Table 3, related to Fig. 6.** Table showing the targeted isoforms of the ASOs that were tested *in vivo*. ASOs highlighted in blue are the ASOs that were selected for further time course studies.

| C1r ASO | C1ra_premRNA | C1rb_premRNA | C1ra_mRNA | C1ra_3UTR |
| --- | --- | --- | --- | --- |
| C1r_1 | Yes | Yes | No | No |
| C1r_2 | Yes | Yes | No | No |
| C1r_3 | Yes | Yes | No | No |
| C1r_4 | Yes | Yes | No | No |
| C1r_8 | Yes | Yes | Yes | Yes |
| C1r_10 | Yes | Yes | Yes | Yes |

| C1s ASO | C1s1_premRNA | C1s2_premRNA | C1s1_mRNA | C1s1_3UTR |
| --- | --- | --- | --- | --- |
| C1s_4 | Yes | No | Yes | No |
| C1s_5 | Yes | Yes | No | No |
| C1s_8 | Yes | No | Yes | Yes |
| C1s_9 | Yes | No | Yes | Yes |
| C1s_10 | Yes | No | Yes | Yes |
| C1s_11 | Yes | Yes | No | No |
| C1s_12 | Yes | No | No | No |
| C1s_13 | Yes | Yes | Yes | Yes |
| C1s_14 | Yes | Yes | Yes | No |

| C4 ASO | C4b_premRNA | C4a_premRNA | C4b_mRNA | C4b_3UTR |
| --- | --- | --- | --- | --- |
| C4_1 | Yes | Yes | Yes | Yes |
| C4_3 | Yes | Yes | No | No |
| C4_5 | Yes | Yes | No | No |
| C4_6 | Yes | Yes | No | No |
| C4_10 | Yes | Yes | Yes | Yes |
| C4_12 | Yes | Yes | No | No |
| C4_14 | Yes | Yes | Yes | No |
| C4_17 | Yes | Yes | Yes | No |
| C4_23 | Yes | Yes | No | No |

To evaluate the time course of target KD, we performed *in vivo* studies with the top ASOs. Brain tissues were collected at 4, 8, and 12 weeks post ICV ASO (100 μg) treatments. Consistent with the 2-week study, all ASOs were well tolerated and did not cause body weight changes compared with saline treated mice (Fig. S7A,B). Brain target gene mRNA expression analysis revealed that ASOs had the strongest KD effect at 4 weeks post dose (Fig. 6E). While C1r_8, C1s_11, and C1s_14 mRNA KD effects began to normalize by 12 weeks, C1r_10 and C1s_8 showed sustained mRNA KD over the 12-week period. Mice that received any of the three C4 ASOs showed sustained KD of C4 mRNA over 12 weeks, with C4_3 being the best (Fig. 6E). We also quantified the C4 protein level in the mouse brain using western blots and found that C4 protein level was well correlated with C4 mRNA level after ASO treatments (Fig. S7C). Based on these results we selected C1r_10, C1s_8 and C4_3 as the best ASOs for an efficacy study in the AD model.

### Targeting the CP with ASOs rescues AD model synapse loss

We next tested the selected C1r, C1s and C4 ASOs, in an efficacy study assessing plaque-associated synapse loss in PS2APP;Thy1-GFP mice after a 100 μg ICV injection of targeting ASOs, a NT control ASO, or a saline control. All of the ASOs were well tolerated by the PS2APP mice at this dose (Fig. S8A,B). Eight weeks after ASO injection, brain C1r, C1s or C4 expression was significantly reduced by their respective ASO treatments in the PS2APP mice compared with saline or NT ASO injected mice. On the other hand, liver C1r, C1s or C4 mRNA levels were unchanged (Fig. 7A). Consistently, brain C4 protein level was also reduced by the C4 ASO, whereas liver C4 protein level was not affected by the C4 ASO (Fig. S8C,D). In addition, the expression of the three inflammatory maker genes (Cst7, Gpnmb, Oasl2) was not changed by any of the ASO treatments in these amyloid mice (Fig. S8E). Consistent with previous studies in PS2APP mice, there was ∼50% reduction in spine density near plaques compared to away from plaque in saline injected mice. The control ASO (NT) did not affect spine density ratio, whereas C1r-targeting, C1s-targeting and C4-targeting ASOs, each significantly improved plaque-associated spine loss (Fig. 7B,C). At the same time, none of the ASOs changed the amyloid plaque density or plaque size (Fig. 7D,E). Comparison of the degree of synapse rescue with the amount of residual mRNA from the experimental results with C1r, C1s and C4 ASO treatment, as well as the homozygous and heterozygous C4 KO study, showed a good correlation between KD and synapse rescue. This is true when KD was evaluated on the mRNA level (Fig. 7F) or on the protein (C4 only) level (Fig. 7G). Taken together, the data suggest that KD of the classical pathway can dose-dependently rescue synapse loss, and targeting the CP with ASOs could be a viable therapeutic approach, with strong knockdown being desirable.

**Fig. 7.**
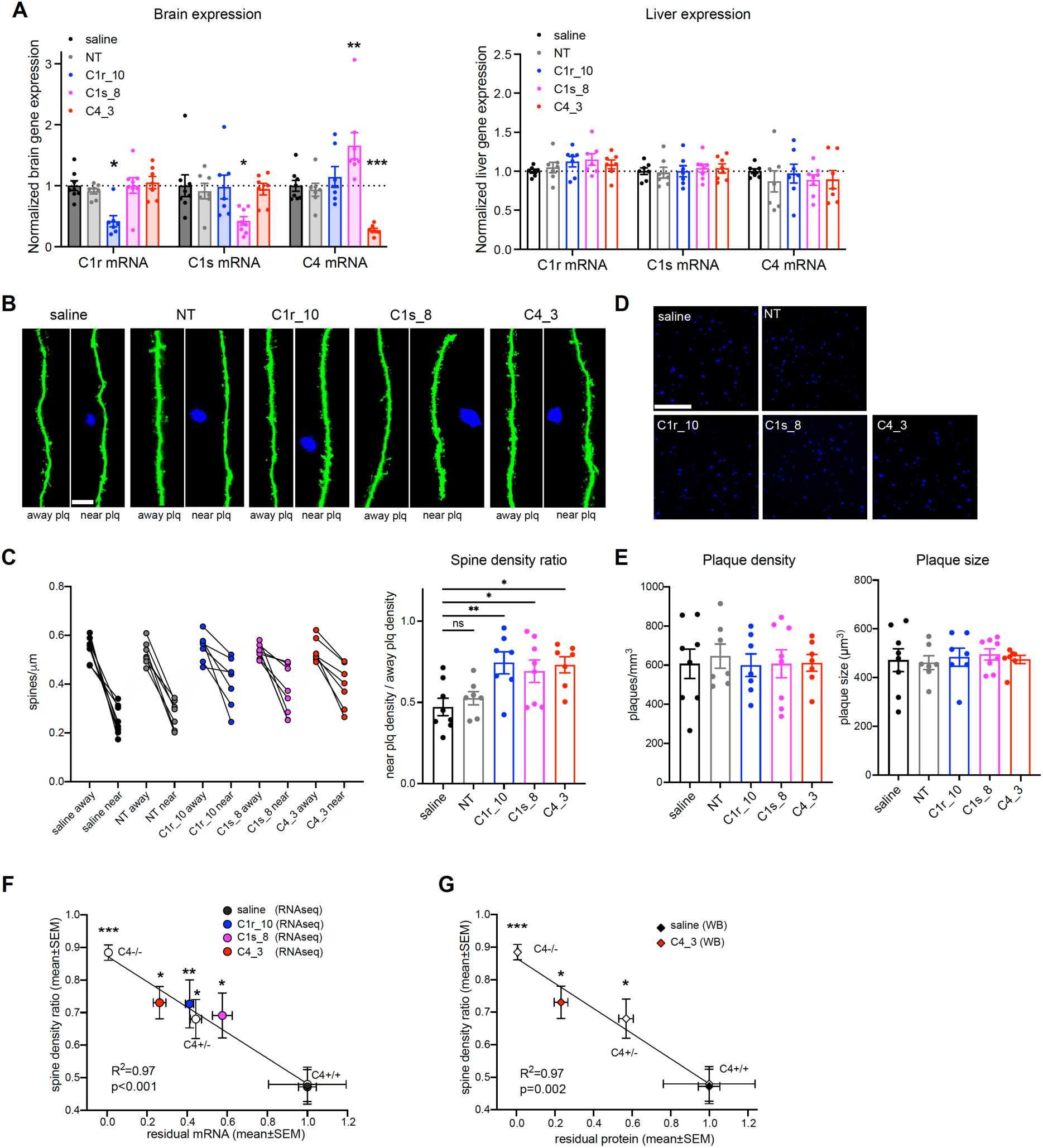
C1r, C1s and C4 ASOs rescues spine loss in PS2APP mice. (**A**) Relative mRNA expression of C1r, C1s or C4 as indicated by the *x-axis* in the mouse brain (*left*) or liver (*right*) two-months after ICV injection of saline, control ASO (NT), C1r ASO, C1s ASO or C4 ASO in PS2APP mice. Each dot represents one mouse, n = 7-8. Data are represented by mean +/- SEM. *, p < 0.05, **, p < 0.01, ***, p < 0.001, compared with saline condition, two-way ANOVA with Tukey’s multiple comparisons test. (**B**) Similar to Fig. 1B, representative images of GFP-expressing neuronal dendrites that are near or away from Aβ plaques in mice injected with saline or corresponding ASOs as indicated. Scale bar, 20 μm. (**C**) Quantification of spine. *Left*, average spine density of dendrites away from plaque and near plaque. *Right*, spine density ratio (average near plaque spine density divided by average away plaque spine density from the same mouse). Each dot represents one mouse (average of 6 dendrites). n = 7-8 mice. Data are shown as mean +/- SEM. *, p< 0.05, **, p < 0.01, one-way ANOVA with Tukey’s multiple comparisons test. (**D**) Representative images of Aβ plaques labeled by methoxy-X04 in PS2APP mouse brains injected with saline or corresponding ASOs as indicated. Scale bar, 200 μm. (**E**) Quantification of Aβ plaque density and size as in (D), n = 7-8 mice. For (C) and (E), data are shown as mean +/- SEM. *, p < 0.05, **, p < 0.01, one-way ANOVA with Dunnett’s multiple comparisons test. (**F**) Relationship of spine density ratio and the relative residual target gene mRNA expression in PS2APP mouse brains with treatments or genotypes indicated. For ASO injected mice, relative brain expression of C1r, C1s or C4 normalized to saline treated mice is shown. For KO mice, the relative level of gene expression relative to wild-type controls is shown (Fig. S1A,C). Expression data are from bulk RNAseq of mouse brain tissues. (**G**) Relationship of spine density ratio with relative C4 protein expression in PS2APP mouse brains is shown (Fig. S1D, Fig. S6C). For (F) and (G), data are represented by mean +/- SEM, n = 4-8 mice.

## Discussion

Our results demonstrate that inhibition of the CP by CNS-delivered ASOs can reduce complement-dependent synapse loss in a mouse model of AD. These experiments show that even partial reduction of C1s, C1r, or C4 can have beneficial effects and suggest that ASOs achieving stronger target KD levels will have even greater benefits. The beneficial effects of ASO treatment are achieved in the absence of any effect on liver complement levels, consistent with activated glial cells being the source of damaging complement in the brain, rather than leakage of circulating complement from the blood.

The translational potential of these results is supported by our analyses of AD patient human genetics and CNS proteomics. While GWAS studies in AD have highlighted the role of microglia, the genetic implication of the complement CP has remained unclear. AD GWAS hits include genes from the complement system such as *CR1* (complement receptor 1) and *CLU* (clusterin / apolipoprotein J / complement lysis inhibitor) (*27, 34*). However, although *CR1* is a complement regulator, the impact of the AD-associated variant on CNS function is not well understood, and clusterin, which has multiple biological roles beyond terminal pathway inhibition, may influence AD risk through mechanisms independent of complement biology. Our analysis of the Finnish population identified an association between a rare rs782726275 C1s variant and AD, building on the previously identified near-genome wide significant association of the common rs3919533 variant with AD, thereby implicating the CP in AD risk. Furthermore, our finding that C1s elevation is the most robust and significant impact of the common rs3919533 C1s variant is consistent with other datasets showing elevated C1s levels in plasma and CSF associated with the risk allele of this variant (*28, 35, 36*). Together these observations suggest a potential link between elevated C1s levels and an increased susceptibility to AD, providing direct genetic evidence implicating the CP in AD.

Our brain proteomics meta-analysis indicates that the CP is specifically elevated in AD patient brains, while the AP, LP, and terminal pathway components are not consistently elevated. This fits with the pathophysiological role of the CP resembling the physiological function in developmental synapse pruning, where C1q targeting of synapses leads to glial-mediated synapse elimination in response to C3b opsonization, without requiring involvement of the AP. LP or terminal pathways (*1, 2*). Our identification of elevated levels of activated C1s in AD patient CSF also supports involvement of the CP in the CNS and is consistent with the previous finding of elevated levels of activated C4, a substrate of activated C1s, in AD patient CSF (*12, 22*). Overall, our results suggest that elevated levels and/or activation of C1s could be associated with AD, and that the CP is specifically elevated in AD brains.

Together our analyses of AD patient genetics and proteomics and the preclinical ASO efficacy studies indicate the translational potential of targeting the complement CP in AD using nucleic acid-based medicine. While we focused on an amyloidosis model, the beneficial effects of complement reduction observed in Tauopathy models (*10–13*) suggest that ASOs targeting the CP could potentially be used as an orthogonal approach to, or in combination with, anti-Aβ therapy for AD patients. Given the implication of the CP in other diseases, such as FTD and Schizophrenia, this approach could potentially have broad application across indications. While our preclinical studies demonstrate promising proof of concept for therapeutically targeting the CP with ASOs, therapeutic ASOs will need to be designed to target primate CP genes and safety profiles of such ASOs will need to be carefully evaluated. While CNS delivery of nucleic acid-based therapeutics to patients currently requires intrathecal injection, ASO and siRNA clinical trials are currently being performed in AD patients, supporting the feasibility of this approach.

## Materials and methods

### Human brain proteomics meta-analysis

Brain tissue proteomics data from Bai et al (*24*), Higginbotham et al (*25*) and Johnson et al (*26*) were downloaded from supplementary tables when available or synapse. Differential abundance statistics were recalculated using *limma*. For each protein, individual log fold changes were combined into a single value using the *rma()* function in the *metafor* package with the DerSimonian-Laird estimator. Individual p-values for a given protein from each study was combined into a single p-value with the sum of logs or Fisher’s method using the *metap* package. Only datasets where the proteins were detected used to combine log fold changes and p-values. The final meta-analysed p-values were corrected using the Bonferroni-Hochberg method.

### Human CSF samples

Cerebrospinal fluid from aged cognitively normal and clinically diagnosed AD subjects were purchased from PrecisionMed (Carlsbad, CA). In compliance with an institutional review board approved protocol for human CSF collection, all donors provided informed consent for the use of these samples in exploratory biomarker studies. Collections from aged cognitively normal subjects were conducted at PrecisionMed, while collections for AD subjects occurred at contracted sites. Lumbar puncture took place in the morning under fasted conditions. The procedure was performed at L3-L4 or L4-L5 using either a 22g Quincke type spinal needle or a 22g Sprotte atraumatic needle, and the CSF was collected by aspiration. If sufficient fluid could not be obtained in this manner, the gravity drip method was utilized. In the event of bleeding, CSF was discarded until blood was no longer visible within the matrix. The remaining CSF (between 10 and 12.5 mL) was collected into a 15 mL polypropylene conical tube, and then centrifuged at 1200 x g for 10 minutes at room temperature. For AD CSF collected at contracted sites, the supernatant was transferred to a 50 mL polypropylene conical tubes, and then immediately frozen in a -80°C freezer. Samples were shipped on dry ice to PrecisionMed, where they were thawed on ice, aliquoted into 1 mL screw-cap polypropylene tubes, and then frozen in a -80°C freezer. For aged cognitively normal CSF collected at PrecisionMed, the CSF supernatant was immediately aliquoted into 1 mL screw-cap polypropylene tubes and then frozen in a -80°C freezer. These samples were thawed on ice at Genentech and then frozen in a -80°C freezer to match freeze-thaw cycles with the AD CSF sample set. Aged cognitively normal subjects (n=10) were between 66-76 years old with MMSE scores between 25-30. AD subjects (n=28) were between 60-88 years old with MMSE scores between 12-27 (Table 1).

### Total and active C1s LC-MS assays

Quantitation of human C1s (total and active) in cerebrospinal fluid (CSF) samples was achieved through direct digestion using LysC (for active C1s, Promega) or Trypsin (for total C1s, Promega) to generate active C1s peptide (IIGGSDADIK) and total C1s peptide (LLEVPEGR). Preparation of standards and quality controls (QCs) ranged from 0.01-100 ug/mL depending on the sample and analyte. To each standard, QC, or sample, LysC or Trypsin digestion buffer and 45mM TCEP were added and subjected to shaking incubation at 60°C for one hour. The samples were cooled, alkylated by the addition of 140mM IAM (final concentration 12 mM), and incubated at room temperature for 30 minutes in darkness, followed by exposure to light for 20 minutes. Digestion was performed with LysC (Promega) or Trypsin (Promega) overnight at 37°C. The digests were then acidified with formic acid and the isotopically labelled peptide standard (IS) was added to each well, followed by LC-MS analysis. The LC-MS employed a Sciex 7500 mass spectrometer, with an accompanying Waters Acquity UPLC, T3 column; MRM (multiple reaction monitoring) was the type of analysis utilized, with specific transitions monitored for each analyte.

### Human genetics analysis

The UK Biobank is a prospective study of over 500,000 participants aged 40–69 years during the recruitment period from 2006–2010. The process of genotyping, imputation, and quality control steps were performed as described previously (*37*). All participants provided informed consent. Participant ancestries were determined according to the pan-UKBB genetic ancestry definitions found in the UKB return dataset 2442. Blood plasma samples from 54,967 UKB participants were analyzed using Olink proteomic profiling, employing the Olink Explore 3072 platform. The process used for participant selection for UKB Proteomics (including the 46,673 randomized participants and the 6,365 selected by consortia), together with the approach to sample handling, has been already exhaustively reported (*28*). We gained access to this data via the UKB application 44257.

The R package “OlinkAnalyze” (version 3.7.0, available at https://CRAN.R-project.org/package=OlinkAnalyze) was employed for differential expression analysis using Welch’s two-sample t-test. In these tests, linear models were constructed for each protein. These models took the risk SNP carriers (characterizing 1 or 2 copies of the risk allele as ’risk allele carriers’ and 0 copy as ’protective allele carriers’) as the primary factor, with protein NPX values as the dependent variables. Covariates included sex, age, and 10 principal components. The Benjamin-Hochberg method was used for multiple testing correction with a false discovery rate. Differentially expressed proteins were defined as assays with false discovery rates less than 0.05 and visualized using the package OlinkAnalyze in R.

FinnGen is a public-private partnership project that aggregates genotype data from Finnish biobanks (https://www.finngen.fi/en). The latest FinnGen release (Data Freeze 10) consists of over 412,000 samples after quality control with population outliers excluded via principal component analysis based on genetic data. The samples have been linked with harmonized data from several national healthcare related registries. AD (Alzheimer’s disease, wide definition, N=15,617) cases were identified from hospital discharge and cause of death registries having G30 (International Classification of Diseases (ICD)-10) or 29010 (ICD-8) codes, from Finnish-specific Social Insurance Institute (KELA) reimbursement registry having 307 or G30 (ICD-10) codes, and from medicine purchase registry having N06D (Anatomical Therapeutic Chemical, ATC) code. The same criteria were used as exclusion criteria for controls (N=396,564). Patients and control subjects in FinnGen provided informed consent for biobank research, based on the Finnish Biobank Act. Alternatively, older research cohorts, collected prior the start of FinnGen (in August 2017), were collected based on study-specific consents and later transferred to the Finnish biobanks after approval by Fimea, the National Supervisory Authority for Welfare and Health. Recruitment protocols followed the biobank protocols approved by Fimea. The Coordinating Ethics Committee of the Hospital District of Helsinki and Uusimaa (HUS) approved the FinnGen study protocol Nr HUS/990/2017. The FinnGen study is approved by Finnish Institute for Health and Welfare (permit numbers: THL/2031/6.02.00/2017, THL/1101/5.05.00/2017, THL/341/6.02.00/2018, THL/2222/6.02.00/2018, THL/283/6.02.00/2019, THL/1721/5.05.00/2019, THL/1524/5.05.00/2020, and THL/2364/14.02/2020), Digital and population data service agency (permit numbers: VRK43431/2017-3, VRK/6909/2018-3, VRK/4415/2019-3), the Social Insurance Institution (permit numbers: KELA 58/522/2017, KELA 131/522/2018, KELA 70/522/2019, KELA 98/522/2019, KELA 138/522/2019, KELA 2/522/2020, KELA 16/522/2020 and Statistics Finland (permit numbers: TK-53-1041-17 and TK-53-90-20).

The Biobank Access Decisions for FinnGen samples and data utilized in FinnGen Data Freeze 10 include: THL Biobank BB2017_55, BB2017_111, BB2018_19, BB_2018_34, BB_2018_67, BB2018_71, BB2019_7, BB2019_8, BB2019_26, BB2020_1, Finnish Red Cross Blood Service Biobank 7.12.2017, Helsinki Biobank HUS/359/2017, Auria Biobank AB17-5154, Biobank Borealis of Northern Finland_2017_1013, Biobank of Eastern Finland 1186/2018, Finnish Clinical Biobank Tampere MH0004, Central Finland Biobank 1-2017, and Terveystalo Biobank STB 2018001.

Detailed description of the FinnGen analysis pipeline can be found on the FinnGen website (https://finngen.gitbook.io/documentation/methods/phewas). Briefly, genome statistics were analyzed using Scalable and Accurate Implementation of Generalized mixed model (SAIGE v0.36.3.2), which uses saddle point approximation (SPA) to calibrate unbalanced case-control ratios. The first ten genetic PCs, sex, age, and genotyping batch were used as covariates.

### Antisense oligonucleotides (ASOs)

Antisense oligonucleotides were synthesized as described previously (*38*). Chemical structures are described in Table 4. ASOs all have phosphorothioate backbones. In base sequence, “E” corresponds to 5-methyl-C. “F” corresponds to PPG nucleotide. In sugar sequence, “L” indicates LNA, “D” indicates DNA.

**Table 4.**
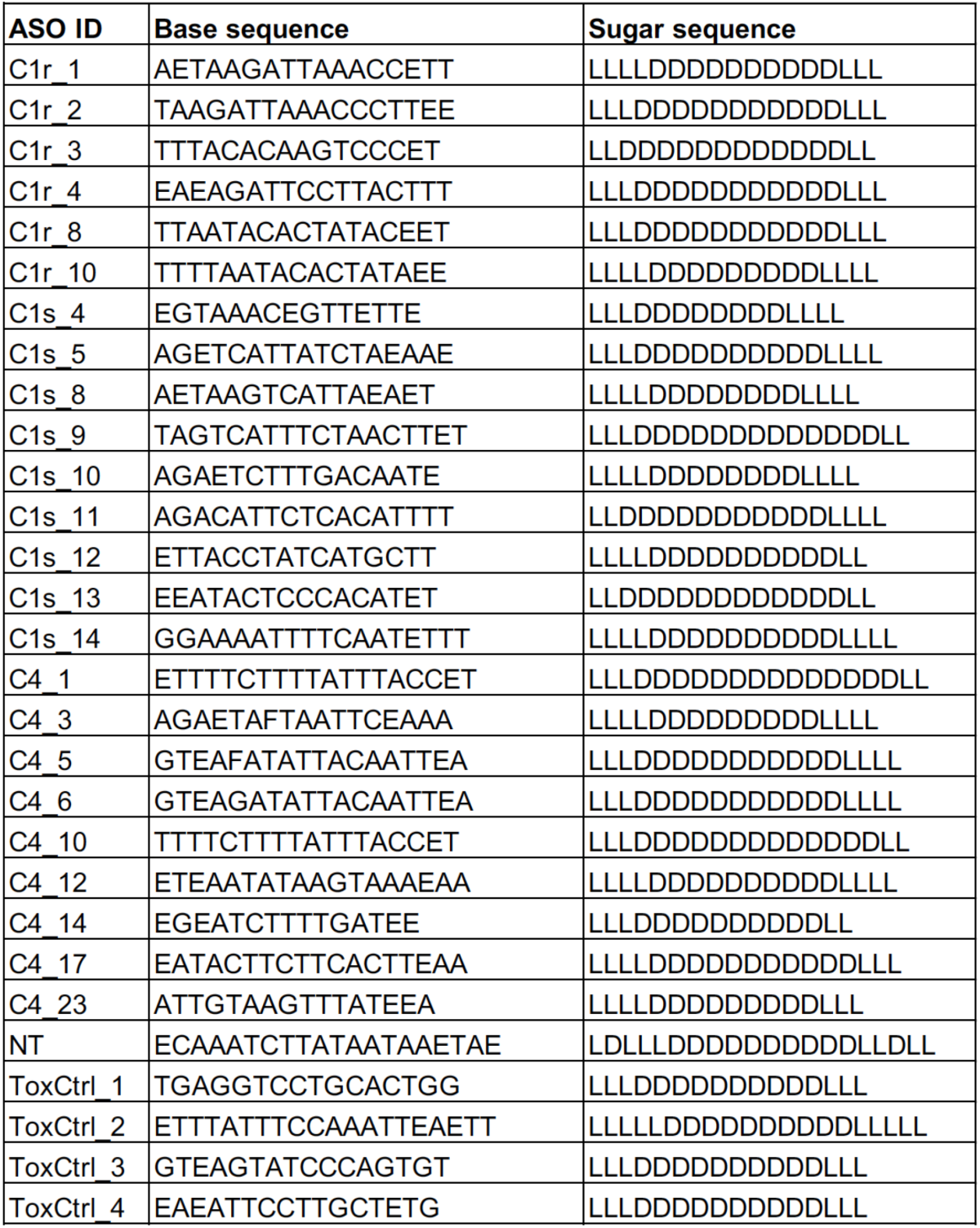
List of sequences of ASOs used in in vivo studies.

### Primary mouse hepatocyte culturing and ASO treatment

PMH cells were purchased from Minerva Imaging. The cells were assessed for cell viability and cultured at 37C, 5 % Co2 and 95% humidity. 25000 cells per well were seeded in 180 ul of WME cell culture medium (WME (Sigma #W1878), 1xPen/Strep/Glutamine (Sigma P4333), 10% (v/v) FBS (Sigma F7524)) in a Collagen coated 96 well cell culture plate (Corning 356407) and left to attach for 24 hours in the incubator. Oligos diluted in PBS (20 μL) were added to the cell cultures from pre-made 96-well dilution plates. The cell culture plates were incubated for 72 hours in the incubator and subsequently harvested and processed for RNA extraction.

### Primary mouse astrocyte culture and ASO treatment

Primary astrocytes were cultured as described previously(*39*). Briefly, P0-P2 mouse brains were dissected out and the brain tissue was disrupted by trituration using a 10 mL serological pipette in cold DMEM media. The homogenate was spun at 300 g for 5 min. The pellet was resuspended in DMEM media and filtered through a 70 μm cell strainer. Dissociated cells were cultured in Flasks with media containing DMEM, 10%FBS and 1% penicillin/streptomycin. Flasks were rinsed with PBS and fresh media was added after 24 h. After ∼12 days, microglia were shaken off (125rpm for 1hr) and fresh media was added back to the flasks. The flasks were then shaken overnight @ 200 rpm and rinsed with PBS twice. Astrocytes were trypsinized, collected and plated at 25K/well in 96-well plate. Cells were allowed to attach and recover for 2 days. Cultures were then incubated with ASOs for 72 hours and cells lysates were collected for RNA extraction. For C4 ASO treatments, only C4 ASOs were added. For C1r and C1s treatments, cytokines were added together with ASOs to stimulate astrocytes in order to get higher C1r and C1s expression levels. Cytokines are: 30 ng/mL TNFα (R&D systems; 410-MT-025/CF) + 3 ng/mL IL-1α (R&D systems; 400-ML-025/CF) + 30 ng/mL IFNγ (R&D systems; 485-MI-100/CF).

### RT-qPCR analysis of target mRNA in primary mouse hepatocytes and primary mouse astrocytes

RNA was extracted using RNAeasy mini kit (Qiagen) from cell lysates and RNA was heat shocked at 90°C for 40 s to melt RNA:LNA duplexes. RT-qPCR was performed using qScript XLT One-Step RT-qPCR ToughMix, Low ROX^TM^ (QuantaBio) following the manufacture’s protocol. RT-qPCR was done on Applied Biosystems ViiA 7 or QuantStudio 7 Flex Real-Time PCR System. Gene expression assays used for mouse hepatocytes and mouse astrocytes (Thermofisher Scientific or Integrated DNA technologies (IDT)) are listed in Table 5.

**Table 5.** List of gene expression assays used for RT-qPCR.

|  |  |  |  |  |
| --- | --- | --- | --- | --- |
| C4b | ThermoFisher | Mm00437890_m1 | Gene Of Interest | hepatocytes |
| C1s | IDT | Mm.PT.58.8983990 | Gene Of Interest | hepatocytes |
| C1ra | IDT | Mm.PT.58.13565159 | Gene Of Interest | hepatocytes |
| Rplp0 | IDT | Mm.PT.58.43894205 | Housekeeping | hepatocytes |
| C4b | ThermoFisher | Mm00437893_g1 | Gene Of Interest | astrocytes/brain/liver |
| C1ra | ThermoFisher | Mm04206253_g1 | Gene Of Interest | astrocytes/brain/liver |
| C1s1 | ThermoFisher | Mm00663210_mH | Gene Of Interest | astrocytes/brain/liver |
| GAPDH | ThermoFisher | Mm99999915_g1 | Housekeeping | astrocytes/brain/liver |
| C1qa | ThermoFisher | Mm07295529_m1 | Gene Of Interest | brain/liver |
| C1qb | ThermoFisher | Mm01179619_m1 | Gene Of Interest | brain/liver |
| C1qc | ThermoFisher | Mm00776126_m1 | Gene Of Interest | brain/liver |
| C2 | ThermoFisher | Mm00442726_m1 | Gene Of Interest | brain/liver |
| Cst7 | ThermoFisher | Mm00438351_m1 | Gene Of Interest | brain/liver |
| Gpnmb | ThermoFisher | Mm01328587_m1 | Gene Of Interest | brain/liver |
| Oasl2 | ThermoFisher | Mm01201449_m1 | Gene Of Interest | brain/liver |

### Caspase activation assay

3T3-L1 cells were cultivated at approximately 70% confluence in growth media (DMEM (Sigma D0819), 10% BCS (Sigma 12133C), 1mM Sodium Pyrovate (Sigma S8636), 25 ug/ml Gentamicin (Sigma G1397)). Cells were detached with 0.25% Trypsin-EDTA solution (Sigma T4049) and seeded into ViewPlate-96 white (Perkin Elmer, 6005181) at a density of 4000 cells/well. Twenty-four hours post-seeding, 3T3-L1 cells were transiently transfected with Lipofectamine 2000 (Life Technologies, no. 11668019) using 100 nM oligonucleotides dissolved in Opti-MEM (Gibco, no. 31985). Caspase 3/7 activity was determined using the Caspase-Glo 3/7 Assay (Promega, Madison, WI, USA). Reconstituted Caspase-Glo 3/7 reagent was added to the cells 24 h post-transfection, incubated for 60 min, and a back seal (Perkin Elmer, 6005199) was placed at the bottom of the cell plate before luminescence was determined on an EnSight Multimode Plate Reader (PerkinElmer) according to the manufacturer’s instructions. Relative caspase activity was calculated by dividing each well with the median of mock treated wells for the individual plate. All experiments were performed in triplicate. ASOs with relative caspase activity lesser than 20% of the median relative caspase activity of the wells treated with toxic ASOs (ToxCtrl1-4 in Table 4) were labeled as “low caspase”, remaining were labeled as “high caspase”.

### Animals

3-month-old C57BL/6J male mice from Jackson Laboratory (Jax 000664) were used for in vivo ASO screening and time-course experiments. PS2APP mice express human APP with the K670N/M671L mutation and human presenilin 2 with the N141I mutation, driven by the Thy1 and PrP promoter, respectively (*40*). PS2APP mice were crossed to the Thy1-GFP-M line transgenic mice, which express EGFP in a subset of neurons (*41*). 4-6 month-old male GFP-hemizygous and APP-homozygous mice were used for spine density experiments. C1qC KO mice and C4b KO mice were from Jackson laboratory (Jax 029409 and Jax 003643, respectively). All mice were on a C57BL/6J background.

Animals were maintained in accordance with the Guide for the Care and Use of Laboratory Animals of the National Institutes of Health. Genentech is an AAALAC-accredited facility and care and handling procedures of animals were reviewed and approved by the Genentech Institutional Animal Care and Use Committee (IACUC) and followed the National Institutes of Health guidelines. Mice were group-housed up to five mice per cage in individually ventilated cages within animal rooms maintained on a regular light/dark cycle (14:10 h) with *ad libitum* access to food (LabDiet 5010) and water. Animal rooms were temperature and humidity-controlled, between 20-26°C and 30-70%, respectively, with 10-15 room air exchanges per hour. All injections and tissue collections were conducted during the light phase.

### Intracerebroventricular (ICV) injections

Injections (ICV) were performed using a freehand method as described previously (*30*). In brief, animals were anesthetized with isoflurane, the skull was shaved, and the skin was cleaned with ChloraPrep swabs (260100, BD Biosciences). Artificial tears lubricant ophthalmic ointment was applied to the eyes and Buprenorphine SR was given to reduce any pain caused by the procedure. A skin incision was made between the occiput and forehead to identify bregma. Injection coordinates were 1-2 mm to the right of the midline and posterior -0.5 mm from bregma. The needle, fitted with cut polyethylene tubing so that no more than 3 mm of needle was available, was inserted through the skull to a depth of -2.8 mm and 30 sec was given to allow for the brain to seal around the needle. A bolus 5 μL injection was given at a rate of 0.5 μL/s, using a 26-gauge 25 μL Hamilton syringe. One min after injection the needle was withdrawn. The incision was closed with VetBond tissue glue, and animals were returned to their home cage on a heat pad to recover.

### Mouse observations post-ICV dosing

Acute side effects scoring was performed at 30 min and 1 h post ICV injection. Acute side effects observations include reduced movement, ataxia, and/or hunched body. These observations were scored as follows: 0, normal; 1, mild; 2, moderate; 3, severe. An even more severe score of 4 is given if seizures, death, or catatonic behavior is observed with no improvement after 2 h. Individual body weights were measured pre-ICV injection as well as every 2 weeks post-injection.

### Tissue collection

For ASO screening and time-course studies, mice were anesthetized with isoflurane and decapitated. For KO and efficacy studies, mice were anesthetized with 2.5% tribromoethanol (0.5 mL/25 g body weight) and transcardially perfused with cold PBS. A 2 mm slice through the right hemibrain was collected just anterior to the injection site and stored at 4°C in RNA*later* (Sigma) solution for RNA extraction and qPCR. A second 2 mm slice through the right hemibrain was collected just posterior to the injection site, the tissue was weighed and frozen at -80°C for WB analysis. The left brain hemisphere was drop-fixed in 4% PFA for 48 h and stored in PBS at 4°C for spine density analysis. Samples of liver tissue were also collected.

### RNA extraction from brain and liver and qPCR analysis

Tissues were transferred from RNA*later* to RLT Buffer (Qiagen) lysis reagent and lysed using a TissueLyser (Qiagen). RNA was extracted using the RNeasy Mini QIAcube Kit (74116, QIAGEN) with a QIAcube workstation. RNA was heat shocked at 90°C for 40 s to melt RNA:LNA duplexes and qPCR was performed on an Applied Biosystems ViiA7 or QuantStudio 7 Flex thermal cycler using qScript XLT One-Step RT-qPCR ToughMix, Low ROX^TM^ (QuantaBio). TaqMan assay probes were obtained from ThermoFisher Scientific (Table 5).

### Ex vivo imaging analysis of dendritic spine density relative to amyloid plaques

We performed dendritic spine imaging ex vivo from the somatosensory cortex of PS2APP mice that also expressed Thy1-GFP-M transgene (*11*). Animals were injected intraperitoneally with Methoxy-X04 (10 mg/kg) to label Individual plaques 24 hours before tissue collection (*42*). Following tissue collection, hemibrains were embedded in agarose for imaging. We imaged spines on apical dendrites of neurons in somatosensory cortex via a 2-photon microscope (Ultima In Vivo Multiphoton Microscopy System; Bruker) using a Ti:sapphire laser (MaiTai DeepSee Spectra Physics; Newport) tuned to 840 nm and a 60x numerical aperture 1.0 objective lens (Olympus) with pixel resolution of 0.1 μm/pixel across a 1024*1024 pixel field of view (FOV) using 0.5 μm steps. In each mouse, 6 cells (1 dendrite/cell) were collected. We defined “near plaque” as a FOV containing a dendrite that is within 20 μm near a plaque and “away from plaque” as an FOV containing only a dendrite with no visible plaque present in the FOV and at least 100 μm outside of the FOV. A custom, semi-automated image analysis routine in MATLAB (MathWorks) was used to generate dendritic spine density. The density is calculated as the total number of spines divided by the length of the corresponding dendrite. For plaque density measurements, larger image stacks (200 μm depth) were collected using a 20x objective across a 1024*1024 pixel FOV with 2 μm steps over the same region as spine density measurements. Plaque density was quantified by a threshold-based MATLAB routine designed to automatically identify methoxy-X04-labeled plaques. Imaging and analysis were performed under blinded conditions.

### Immunoblotting

Immunoblotting was performed as previously described (*39*). Mouse brain or liver tissues were homogenized in cold RIPA buffer (50 mM Tris-HCl, 150 mM NaCl, 2 mM EDTA, 1% NP-40, 0.1% SDS) supplemented with phosphatase and protease inhibitors, using a TissueLyser (2 × 30 Hz, 3 min at 4°C; QIAGEN). After homogenization, samples were centrifuged at 20,000 g for 20 min and the supernatants were transferred into new tubes. The supernatants were then boiled in reducing SDS sample buffer. The protein samples were separated by Novex Tris-Glycine SDS-PAGE gels (Invitrogen) and transferred to nitrocellulose membranes (Bio-Rad). Membranes were then blocked with Blocking Buffer for Fluorescent Western Blotting (Rockland, MB-070) for one hour (RT) and incubated with primary antibodies (in blocking buffer with 0.1% Triton X-100, 4°C overnight), and then fluorescence-conjugated secondary antibodies (IRDye 800CW and IRDye 680RD; LI-COR biosciences) (two hours RT). Fluorescence signals were detected by Odyssey imager (LI-COR biosciences). Data analysis was done using Image J. Primary antibodies used for western blots are: anti-C4 (Abcam, ab11863), anti-α-Tubulin (Sigma, T5168), anti-GAPDH (Cell Signaling Technology, #5174).

### C1q protein analysis

C1q protein quantification assay was developed in-house based on Gyrolab immunoassay platform using Gyrolab Bioaffy CDs. The capture antibody was anti-C1q (JL-1), biotin conjugated monoclonal antibody. The detection antibody was anti-C1q (JL-1), Alexa 647 conjugated. Mouse brains were homogenized using RIPA buffer as described above and supernatants of the brain lysates were used for C1q quantification.

### Statistical analysis

Data were analyzed using GraphPad Prism. Statistical testing used for each analysis was listed in the corresponding figure legend. Data are represented by mean +/- SEM. Comparisons were considered statistically significant when p < 0.05.

## List of Supplementary Materials

Fig. S1, related to Fig. 1B.

Fig. S2, related to Fig. 1D.

Fig. S3. C1q and C4 het mice have ∼ 50% reduction of mRNA and protein in the brain.

Fig. S4. Expression of C1r, C1s, C2 and C4 isoforms in mice.

Fig. S5, related to Fig. 5.

Fig. S6, related to Fig. 6. Additional toxicity and KD measures for single time point *in vivo* ASO studies.

Fig. S7, related to Fig. 6. Additional toxicity and KD measures for *in vivo* ASO time course studies.

Fig. S8, related to Fig. 7, toxicity analysis and C4 protein measurement in ASO-treated PS2APP mice.

## Acknowledgements

We want to acknowledge the participants and investigators of the FinnGen study. We would like to thank Jianhui Zhu and John Chen from NovaBioAssays for MS assay support.

## Author contributions

Y.W., T.W., A.G., M.C.T, L.K., A.M., J.C.T., C.W., S.S., W.J.M, J. E., A. E., H. M. G., and J.E.H. contributed to the design of the work and interpretation of data. Y.W., T.W., A.G., M.C.T, A.M., S.P., L.S., C.W., S.B., A.M contributed to the acquisition and analysis of data. J.E.H conceptualized the work. Y.W. and J.E.H wrote the manuscript. All authors read and edited the manuscript.

## Data and materials availability

All data are available in the main text or the supplementary materials.

