## Supplementary Figures (S1-S8) for "Therapeutically targeting the classical complement pathway with antisense oligonucleotides in Alzheimer’s disease"

Fig. S1

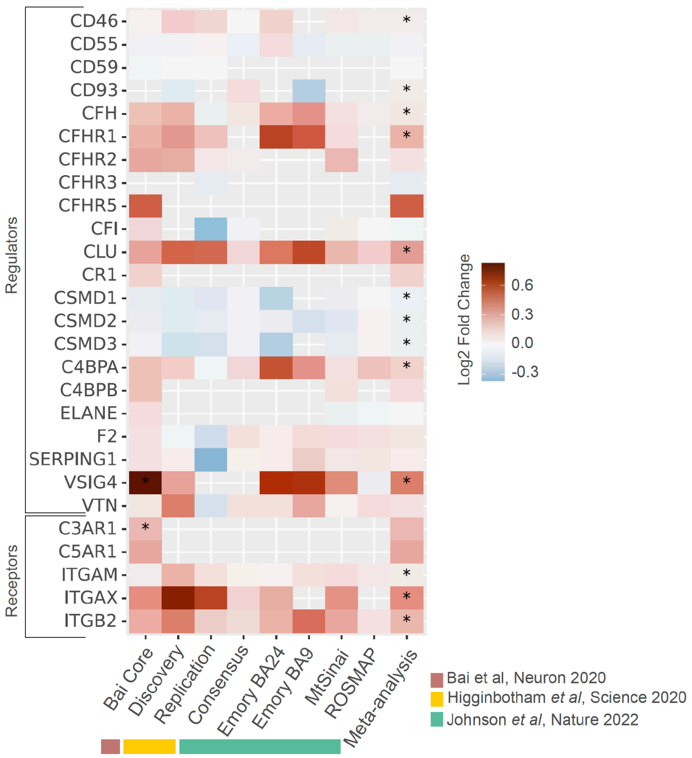


**Fig. S1, related to Fig. 1B**

Heatmaps for complement protein regulators and receptors.

Fig. S2

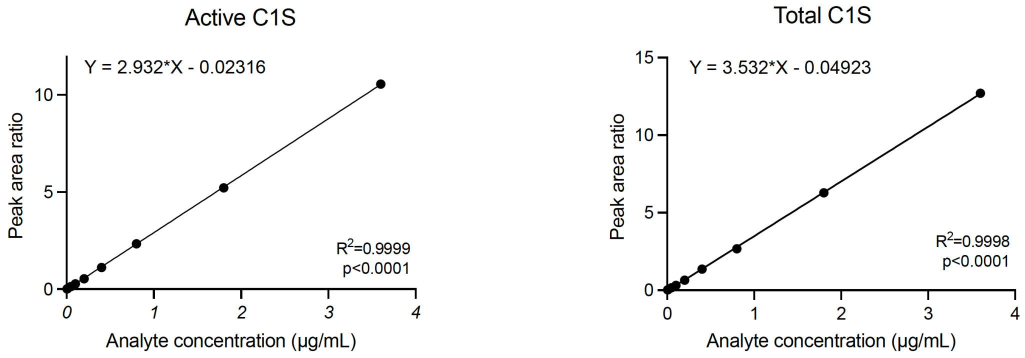


**Fig. S2, related to Fig. 1D**

Mass spectrometry signal calibration curves in CSF showing linearity of quantitation of isotopic heavy-labeled signature peptides for active C1s (IIGGSDADIK) upon LysC digestion and total C1s (LLEVPEGR) upon trypsin digestion. Both active and total C1s assays were qualified for accuracy and precision using recombinant standard spike-ins in CSF at a concentration range of 0.005 – 3.6 $\mu$g/mL showing accuracy and precision within 10%.

Fig. S3

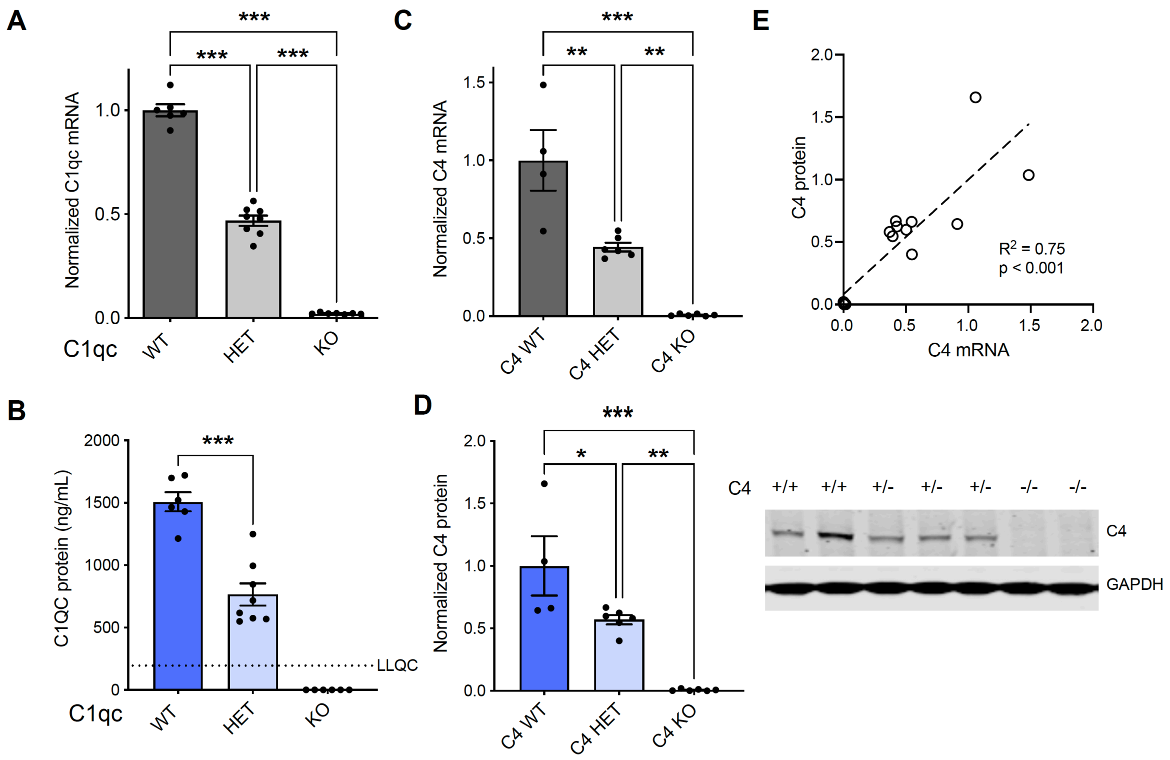


**Fig. S3. C1q and C4 het mice have ~ 50% reduction of mRNA and protein in the brain**

(**A**) Relative expression of C1qc mRNA in the brains of mice with genotypes as indicated, measured by RT-qPCR. C1qc expression was normalized to the mRNA level in WT mice after normalization to GAPDH. (**B**) C1qc protein levels measured by ELISA in the brains of mice with genotypes as indicated. (**C**) Relative expression of C4 mRNA in the brains of mice with genotypes as indicated, measured by RT-qPCR. C4 expression was normalized to the mRNA level in WT mice after normalization to GAPDH. (**D**) Representative immunoblots (*right*) and quantification (*left*) for C4 protein in brain lysates from C4 WT, HET and KO mice. Protein levels were normalized to C4 protein level in WT mice. (**E**) Correlation of C4 mRNA level with C4 protein level in the same mouse. Each dot represents one mouse, n = 4-8. Data are represented by mean +/- SEM, *, $p < 0.05$, ***, $p < 0.001$, one-way ANOVA with Tukey's multiple comparisons test.

Fig. S4

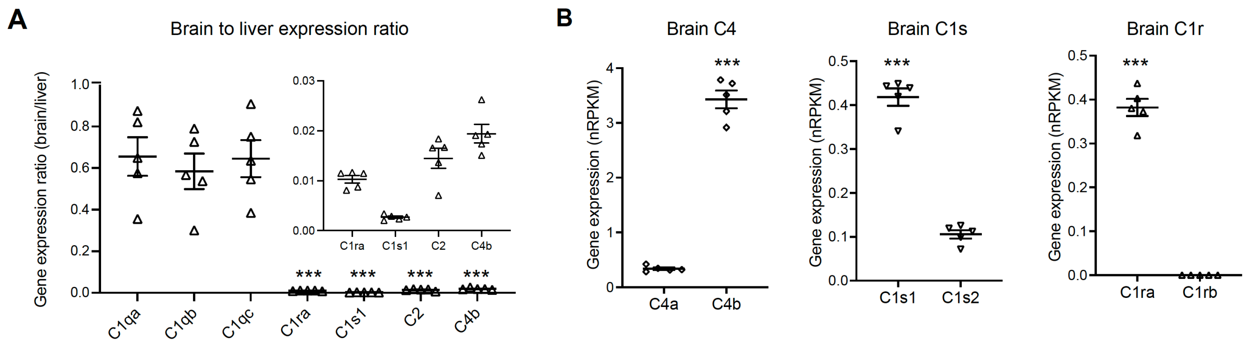


**Fig. S4. Expression of C1r, C1s, C2 and C4 isoforms in mice.**

(**A**) Relative expression of C1q, C1r, C1s, C2 and C4 in the mouse brain compared to that in the liver measured by RT-qPCR. Inset shows C1r, C1s, C2 and C4 expression ratios with zoomed in y-axis values. Each dot represents a mouse, n = 5. C1r, C1s, C2 and C4 expression ratios were statistically significantly lower compared with C1qa, C1qb, or C1qc expression ratio, ***, $p < 0.001$, one-way ANOVA with Dunnett's multiple comparisons test. (**B**) Gene expression of C1r, C1s, and C4 isoforms in mouse brain measure by bulk RNAseq (GSE196401). n =5 mice. ***, $p < 0.001$, Student's t-test. Data are represented by mean +/- SEM.

Fig. S5

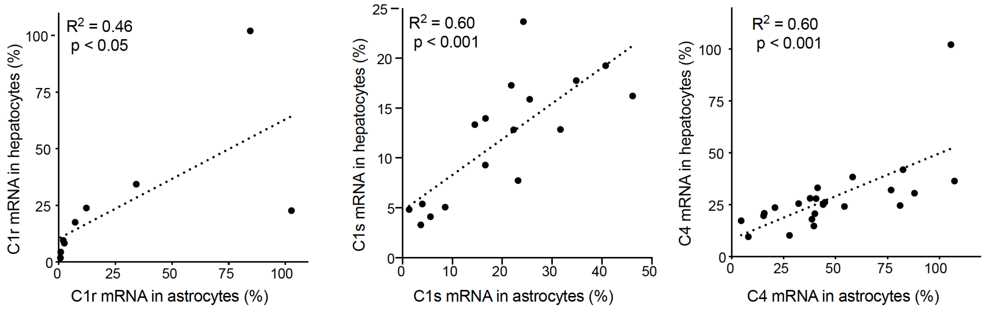


**Fig. S5, related to Fig. 5.**

Correlation of knockdown of C1r (*left*), C1s (*middle*) or C4 (*right*) by the respective ASOs in primary hepatocytes and primary astrocytes. Each dot represents one ASO. All ASOs that were selected for secondary astrocyte screen are shown in the graphs.

Fig. S6

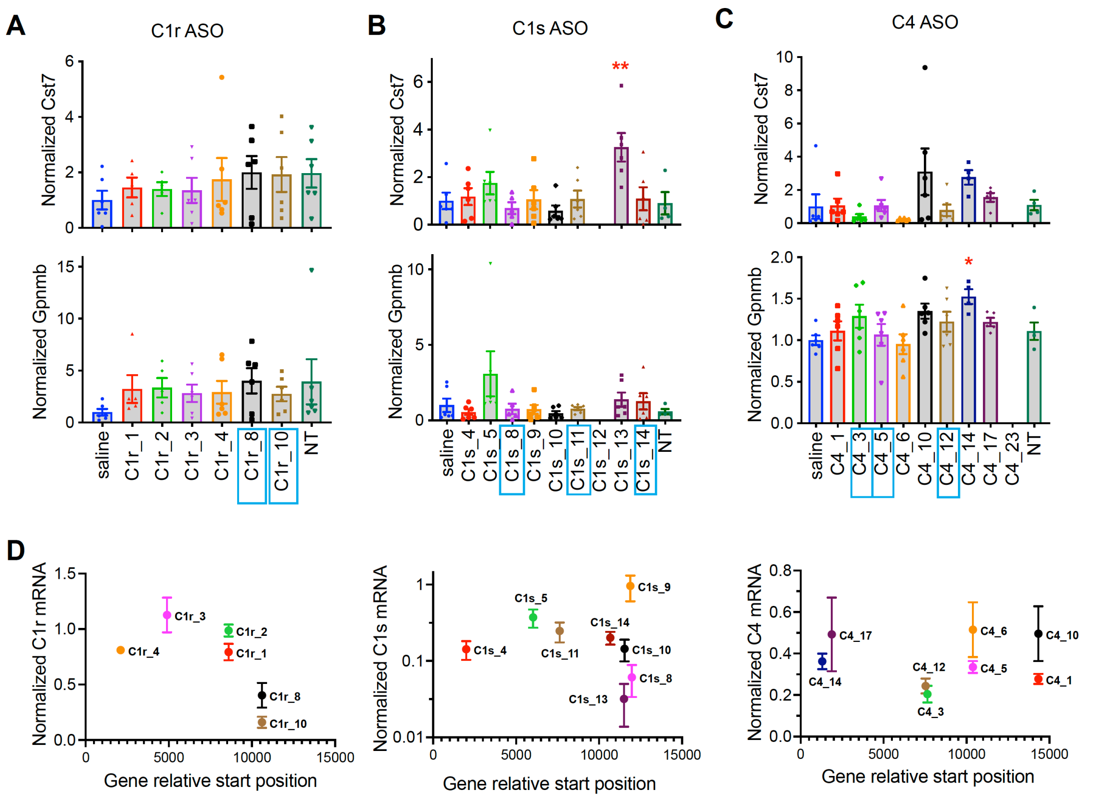


**Fig. S6, related to Fig. 6. Additional toxicity and KD measures for single time point *in vivo* ASO studies**

(**A-C**) Related to Fig. 6A-C, relative inflammation marker genes Cst7 (top) and Gpnmb (bottom) expression measured by RT-qPCR from mouse brain tissues collected 14 days after ICV injection of ASOs as indicated. Data are normalized to Cst7 or Gpnmb expression in saline injected mice respectively. Each dot represents a mouse, n = 4-6 mice per condition. Data are represented by mean +/- SEM. *, $p < 0.05$, **, $p < 0.01$, one-way ANOVA with Dunnett's multiple comparisons test; compared with saline condition. (**D**) X-axis illustrates the binding position on the chromosome of each ASO as indicated, and y-axis shows the relative expression of target gene mRNA in mouse brains after 14-day ASO treatments as indicated. mRNA expression data are represented by mean +/- SEM.

Fig. S7

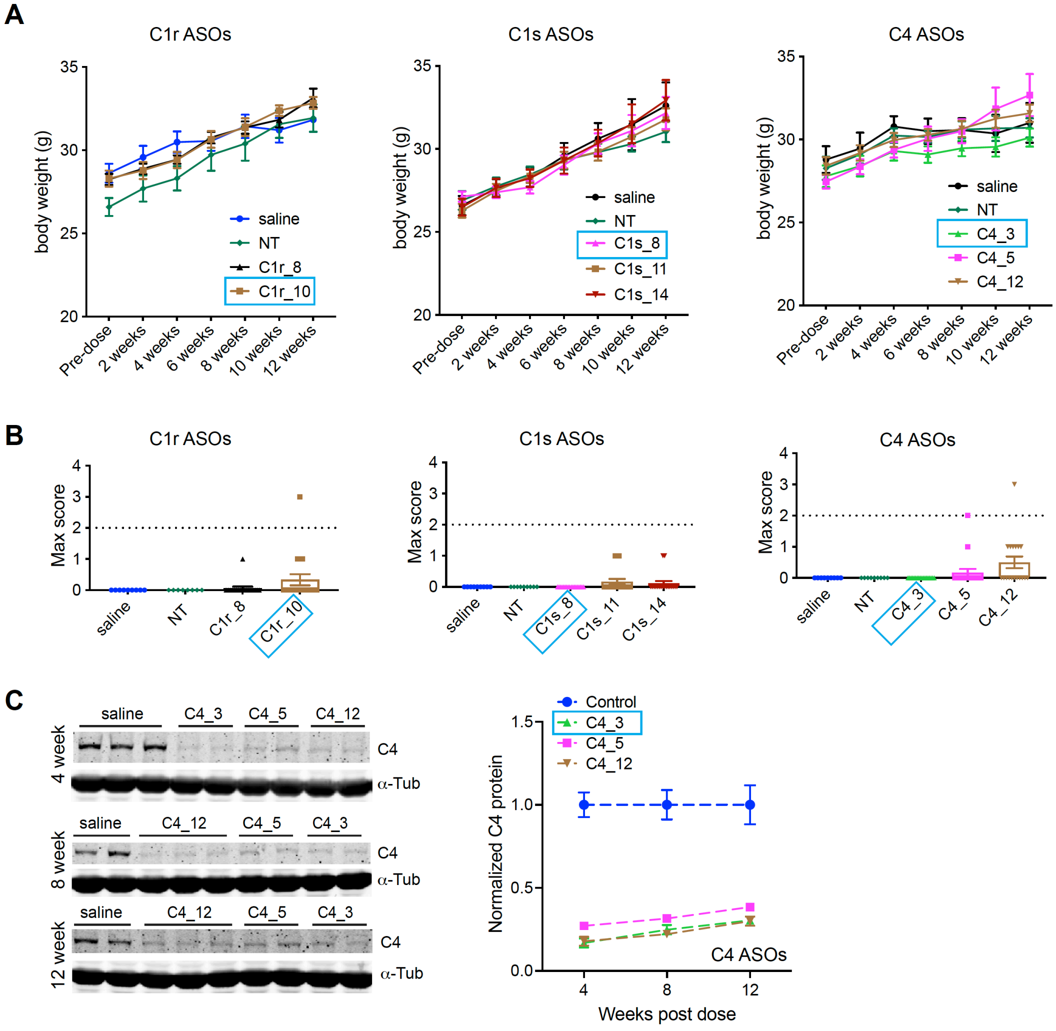


**Fig. S7, related to Fig. 6. Additional toxicity and KD measures for *in vivo* ASO time course studies**

(**A**) Related to Fig. 6E, mouse body weights before and every two-week after saline or ASO injection as indicated. Data are represented by average mouse body weights per condition +/- SEM. (**B**) Related to Fig. 6E, acute toxicity scores after ICV injection of ASOs as indicated. Data are represented by mean +/- SEM, n = 9-18 mice per condition. Note the large n number for the acute toxicity measurements is because the mice were taken down gradually at different time points. (**C**) Representative immunoblots for C4 and α-tubulin (loading control) in mouse brain tissue lysates collected 4-week, 8-week and 12-week after C4 ASO injections as indicated. Quantification of C4 protein level from blots shown (left) are shown on the right.

C4 protein levels were first normalized to α-tubulin, and then the normalized values were plotted relative to the average values from control samples (saline and NT ASO injected samples), n = 5-6 mice per condition. Data are represented by mean +/- SEM.

Fig. S8

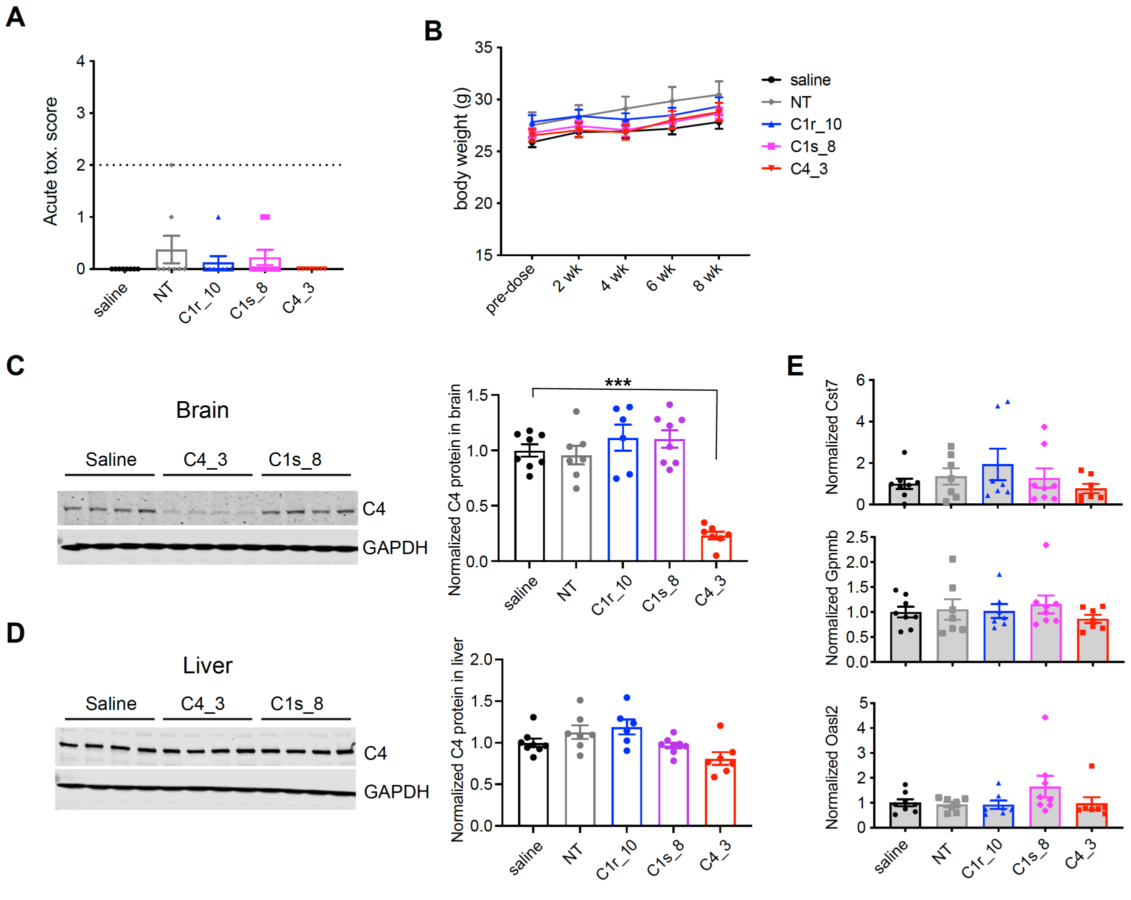


**Fig. S8, related to Fig. 7, toxicity analysis and C4 protein measurement in ASO-treated PS2APP mice**

(**A**) Similar to Fig. 6A, acute ASO induced toxicity after ICV injection was assessed and mouse behavioral toxicity responses were scored (see methods). (**B**) Mouse body weights before and after saline/ASO injection. Data are represented by average mouse body weights per condition +/- SEM, n = 7-8 mice per condition. (**C**) Representative immunoblots (left) and quantification (right) for C4 protein in brain tissue lysates from mice injected with saline or ASOs as indicated. (**D**) Representative immunoblots (left) and quantification (right) for C4 protein in liver tissue lysates from PS2APP mice injected with saline or ASOs as indicated. (**E**) RT-qPCR measurements of relative expression of inflammation marker genes Cst7, Gpnmb and Oasl2 in PS2APP mouse brains injected with saline or ASOs as indicated. Gene expression was normalized to its expression in saline treated mice respectively after normalization to GAPDH. Each dot represents one mouse, n = 7-8. Data are represented by mean +/- SEM. ***, $p < 0.001$, compared with saline condition, one-way ANOVA with Dunnett's multiple comparisons test.
